# Translocation of polyubiquitinated protein substrates by the hexameric Cdc48 ATPase

**DOI:** 10.1101/2021.10.21.465358

**Authors:** Zhejian Ji, Hao Li, Daniele Peterle, Joao A. Paulo, Scott B. Ficarro, Thomas E. Wales, Jarrod A. Marto, Steven P. Gygi, John R. Engen, Tom A. Rapoport

## Abstract

The hexameric Cdc48 ATPase (p97 or VCP in mammals) cooperates with its cofactor Ufd1/Npl4 to extract polyubiquitinated proteins from membranes or macromolecular complexes for degradation by the proteasome. Here, we clarify how the Cdc48 complex unfolds its substrates and translocates polypeptides with branchpoints. The Cdc48 complex recognizes primarily polyubiquitin chains, rather than the attached substrate. Cdc48 and Ufd1/Npl4 cooperatively bind the polyubiquitin chain, resulting in the unfolding of one ubiquitin molecule (initiator). Next, the ATPase pulls on the initiator ubiquitin and moves all ubiquitin molecules linked to its C-terminus through the central pore of the hexameric double-ring, causing transient ubiquitin unfolding. When the ATPase reaches the isopeptide bond of the substrate, it can translocate and unfold both N- and C-terminal segments. Ubiquitins linked to the branchpoint of the initiator dissociate from Ufd1/Npl4 and move outside the central pore, resulting in the release of unfolded, polyubiquitinated substrate from Cdc48.

## INTRODUCTION

The Cdc48 ATPase in *Saccharomyces cerevisiae* and its p97 (VCP) orthologs in metazoans extract polyubiquitinated substrate polypeptides from membranes or macromolecule complexes and generally deliver them to the proteasome for degradation (Bodnar and Rapoport, 2017a; van den Boom and Meyer, 2018). For example, in endoplasmic reticulum (ER)-associated protein degradation (ERAD), Cdc48/p97 pulls misfolded, polyubiquitinated proteins out of the ER membrane for their subsequent degradation (Wu and Rapoport, 2018). Cdc48/p97 consists of an N-terminal N domain and two ATPase domains (D1 and D2) (Bodnar and Rapoport, 2017a) (**Figure S1A**). Six molecules of the ATPase form a double-ring structure with a central pore. In ERAD and many other processes, Cdc48/p97 cooperates with the heterodimeric Ufd1/Npl4 (UN) cofactor. Like Cdc48/p97, UN is found in all eukaryotic cells and is essential for their viability. UN recruits substrates to the Cdc48/p97 ATPase by interacting with the attached K48-linked polyubiquitin chain. Subsequently, the ATPase uses ATP hydrolysis to translocate the polypeptide through the central pore, thereby causing its unfolding (Blythe et al., 2017; Bodnar and Rapoport, 2017b). The critical role of human p97 in protein quality control is highlighted by mutations that cause neurodegenerative proteopathies (Johnson et al., 2010; Kimonis et al., 2008; Watts et al., 2004). p97 is also an important cancer drug target, as inhibitors suppress the proliferation of multiple tumors (Anderson et al., 2015; Le Moigne et al., 2017). Despite its importance, the mechanism by which Cdc48/p97 processes its substrates remains poorly understood.

In previous work, we determined cryo-EM structures of Cdc48 in complex with the UN cofactor and a polyubiquitinated model substrate (Twomey et al., 2019) (**Figure S1A**). The structures represented an initiation state of substrate processing prior to ATP hydrolysis. The D1 and D2 domains formed stacked hexameric rings, while Npl4 formed a tower-like structure above the D1 ring (**Figure S1A**). Most of the Ufd1 molecule was invisible. Three consecutive ubiquitin molecules of the substrate-attached polyubiquitin chain were seen. Two folded ubiquitin molecules (Ub1 and Ub2) were bound to the top of the Npl4 tower, and one ubiquitin molecule was unfolded (the “initiator ubiquitin”), with its N-terminal segment traversing the central pores of both ATPase rings and a subsequent segment bound to a groove in Npl4 (**Figure S1A**). We hypothesized that Cdc48 begins polypeptide translocation by pulling on the N-terminal segment of the initiator ubiquitin, because this segment interacts with the D2 pore loops and the D2 ATPases are responsible for polypeptide movement (Bodnar and Rapoport, 2017b).

Substrate processing by the Cdc48 complex can be divided into three phases--substrate recruitment, translocation, and release-- which are all poorly understood. Our previous in vitro experiments suggested that a polyubiquitin chain is sufficient to initiate translocation (Bodnar and Rapoport, 2017b), while the actual substrate to which the chain is attached plays no role. However, it remained unclear whether in vivo the ATPase complex indeed indiscriminately processes all polyubiquitinated proteins, particularly because this would raise the possibility that all such substrates are unfolded by the ATPase, and that Cdc48/p97 and the 26S proteasome compete with one another for polyubiquitinated proteins. How the ubiquitin chain is recognized by the Cdc48 complex is also unclear. The most mysterious aspect concerns the unfolding of the initiator ubiquitin molecule (Twomey et al., 2019). Ubiquitin is extremely stable, and yet it can be unfolded by a simple binding reaction, without the need for ATP hydrolysis.

Many aspects of the translocation process also remain unclear. If Cdc48 begins translocation by pulling on the initiator ubiquitin, it would need to translocate all ubiquitin molecules positioned between the initiator and substrate (proximal ubiquitins) before it can unfold the substrate (**Figure S1A**). This implies that the ATPase translocates branched polypeptide chains, as each ubiquitin molecule is linked by an isopeptide bond through its C-terminus to K48 of another ubiquitin molecule or to a Lys residue of the substrate. It is unknown how Cdc48/p97 deals with such branchpoints, i.e. whether it translocates and unfolds both polypeptide branches or only one of them. The ability to process branched polypeptides sets the Cdc48/p97 ATPase apart from most other known ATPases, which translocate only linear polypeptide chains. For example, the 26S proteasome recognizes polyubiquitin chains similarly to the Cdc48/p97 ATPase, but it cleaves them off before translocating and degrading an unbranched substrate polypeptide (Greene et al., 2020). Finally, it is unclear how substrate is released from the Cdc48/p97 complex after completion of translocation, so that it can be transferred to the proteasome.

Here, we show that the Cdc48 ATPase complex indeed shows little substrate specificity, processing the majority of polyubiquitinated substrates in a cell. We clarify the mechanisms of all three phases of substrate processing by showing how the ATPase complex unfolds the initiator ubiquitin, how it translocates polypeptides with branchpoints, and how it releases its polyubiquitinated substrates. Our work provides a comprehensive model for protein unfolding by the Cdc48 ATPase and suggests that the ATPase acts both upstream and downstream of the proteasome.

## RESULTS

### The Cdc48 complex recognizes primarily the polyubiquitin chain

We first tested in *S. cerevisiae* cells whether the Cdc48/UN complex binds equally well to all polyubiquitinated proteins or whether it processes only a subset of them. We used quantitative MS to compare the proteome carrying K48-linked polyubiquitin chains with the proteome associated with the Cdc48/UN complex. The experiments were performed in the absence or presence of the proteasome inhibitor bortezomib in cells lacking the drug exporter Pdr5 (Fleming et al., 2002).

To enrich for proteins bound to the Cdc48/UN complex, we expressed FLAG-tagged Npl4 or Ufd1 (Npl4-FLAG and Ufd1-FLAG, respectively) in yeast cells, and subjected cell lysates to immunoprecipitation (IP) with FLAG antibodies bound to beads (**Figure 1A**). Pulling on UN, rather than Cdc48, avoids the co-purification of other cofactors and their associated substrates. The beads were then treated with an excess of trypsin-resistant tandem ubiquitin-binding entity (TR-TUBE) (Tsuchiya et al., 2017), resulting in the transfer of polyubiquitinated proteins from the Cdc48 complex to TUBE. The samples were subjected to trypsin digestion and the resulting peptides labeled with Tandem Mass Tags (TMT). The use of trypsin-resistant TUBE prevented the interference of abundant TUBE peptides in the subsequent MS analysis. Proteins carrying K48-linked polyubiquitin chains were enriched by incubating cell lysates with biotinylated TUBE-K48 (^Bio^TUBE^K48^), a protein that recognizes specifically K48-linked polyubiquitin chains, followed by incubation with streptavidin beads (K48 IP) (**Figure 1A**). Bound polyubiquitinated proteins were again eluted with TR-TUBE, digested with trypsin, and subjected to TMT labeling. Labeled peptides from all samples were mixed and subjected to tandem MS. For each substrate protein detected, we determined its relative abundance in the FLAG IP versus K48 IP (Cdc48/K48 ratio). With either Npl4-FLAG or Ufd1-FLAG pull-downs, most identified substrate proteins have approximately the same Cdc48/K48 ratio (**Figure 1B; 1C**), consistent with Npl4 and Ufd1 being stoichiometric components of the Cdc48/UN complex. Treatment with bortezomib did not significantly affect the Cdc48/K48 ratios (**Figures 1D; 1E**), despite the large accumulation of polyubiquitinated proteins in bortezomib-treated cells (**Figure S1B**). Averaging Cdc48/K48 ratios for each protein shows that 91% of all identified proteins (795 out of 873) had ratios between 0.3 and 3, i.e, did not drastically differ in their enrichment by pulling on the Cdc48 complex or K48-linked ubiquitin chains (**Figure 1F**). Thus, the ATPase complex recognizes primarily the polyubiquitin chain and has little specificity for the attached substrate.

**Figure 1.**
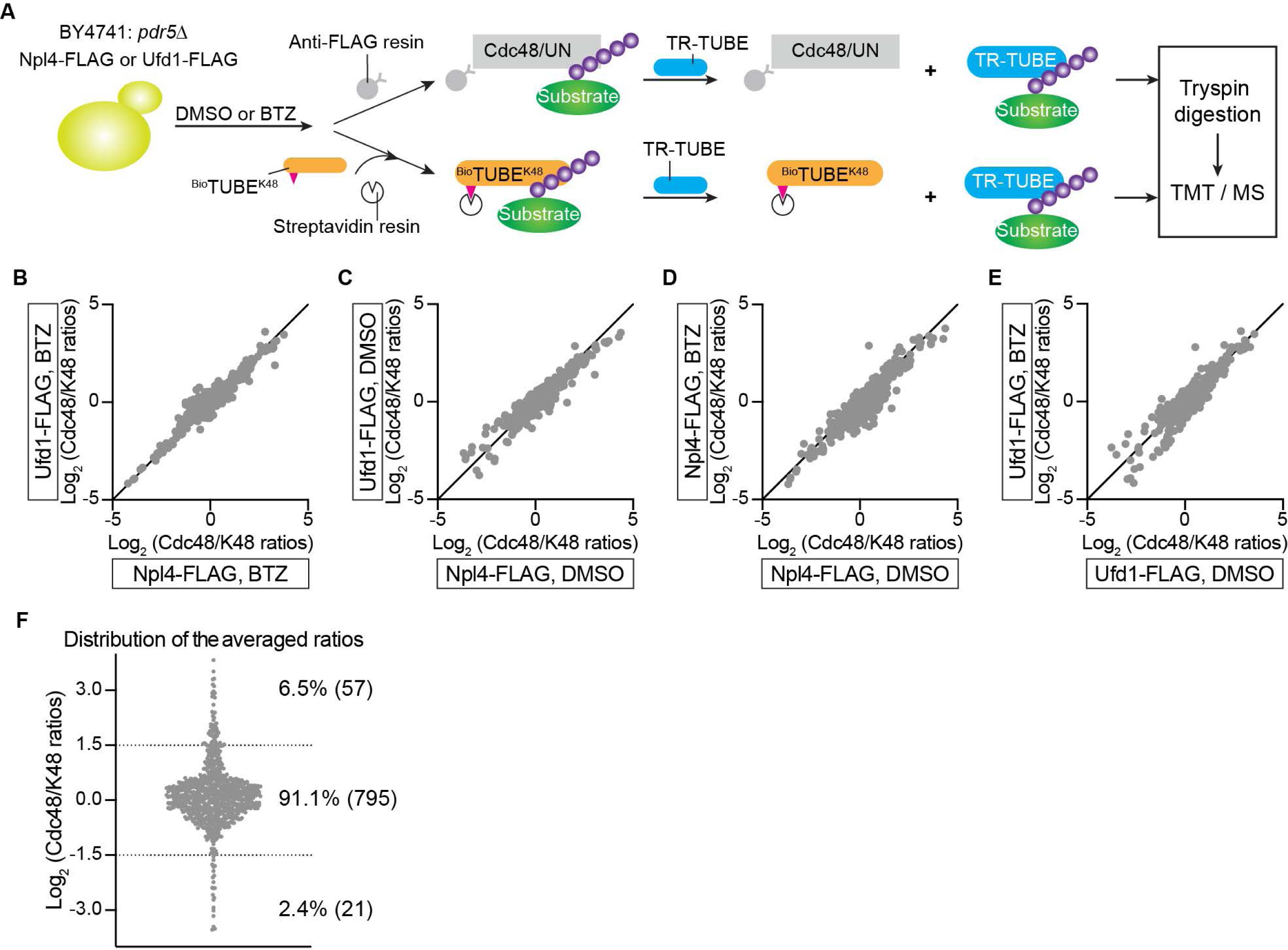
Most proteins carrying K48-linked ubiquitin chains interact with the Cdc48/UN complex. **(A)** *S. cerevisiae* cells lacking the ABC transporter Pdr5 (BY4741:*pdr*5Δ) and expressing Npl4-FLAG or Ufd1-FLAG from plasmids were incubated with or without the proteasome inhibitor bortezomib (BTZ). Cell lysates were incubated with either beads containing FLAG antibodies (Anti-FLAG resin) or biotinylated TUBE recognizing K48-linked ubiquitin chains (^Bio^TUBE^K48^) and streptavidin beads. Polyubiquitinated proteins were eluted with an excess of trypsin-resistant TUBE (TR-TUBE). The samples were subjected to trypsin digestion, TMT labeling, and analysis by tandem MS. **(B)** For each substrate protein detected in BTZ-treated cells, its abundance in the FLAG immunoprecipitation (IP) was divided by its abundance in the K48 ubiquitin pull-down (K48 IP) (Cdc48/K48 ratios). The ratios from Npl4-FLAG and Ufd1-FLAG cells were plotted against each other on a logarithmic scale. **(C)** As in (B), but for untreated cells (DMSO instead of BTZ). **(D)** As in (B), but comparing data from Npl4-FLAG expressing cells with or without BTZ treatment. **(E)** As in (D), but for Ufd1-FLAG expressing cells. **(F)** For each detected protein, the Cdc48/K48 ratios were averaged among all tested cell lysates. Proteins between the dotted lines are enriched or depleted by a factor of less than three.

### Cdc48-facilitated ubiquitin unfolding

We next used in vitro experiments to analyze the molecular mechanism of substrate processing by the Cdc48 ATPase complex, first focusing on the initiation stage, during which a ubiquitin molecule is unfolded without the need for ATP hydrolysis. To follow substrate and ubiquitin unfolding, we performed hydrogen/deuterium exchange (HDX) mass spectrometry (MS) with purified Cdc48 complex and a polyubiquitinated model substrate. The substrate contained a degron sequence derived from the N-end rule pathway (Chau et al., 1989) fused to the fluorescent protein Dendra (Kaberniuk et al., 2017) (**Figure 2A**). The fusion protein was irradiated with UV light, which leads to photoconversion from green to red fluorescence and cleavage of the Dendra polypeptide into two fragments that remain non-covalently associated with one another. The irradiated fusion protein was then incubated with purified ubiquitination enzymes to attach a single chain of 10-25 K48-linked ubiquitin molecules to a Lys residue in the N-terminal segment, resulting in Ub(n)-Dendra (Bodnar and Rapoport, 2017b) (**Figure 2A**).

**Figure 2.**
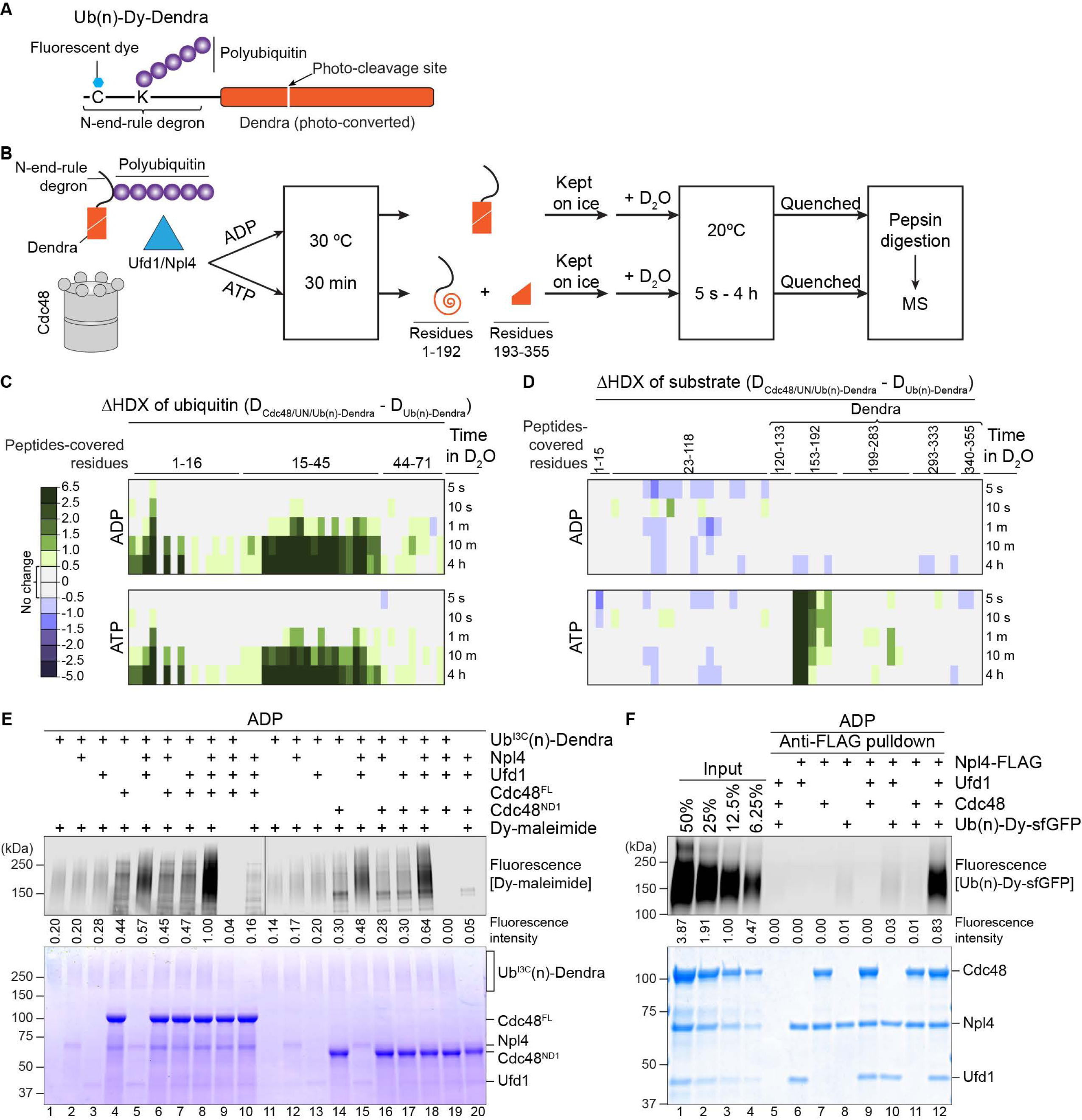
Cdc48-mediated ubiquitin unfolding. **(A)** Scheme of the Ub(n)-Dendra substrate employed for in vitro experiments. **(B)** Experimental protocol of the HDX experiments. **(C)** Photoconverted Ub(n)-Dendra was incubated with or without Cdc48/UN complex in the presence of ADP or ATP. The samples were then subjected to HDX for different time periods, the proteins were subjected to proteolytic cleavage, and the deuteration of ubiquitin peptides determined by MS. Ubiquitin peptides covered the indicated amino acids. Shown is the difference in the deuterium level (in Daltons) caused by binding of the Cdc48 complex (color scale shown in the left panel). **(D)** As in (C), but for the Dendra-fusion protein. Peptides covering amino acids 153-192 contain the two N-terminal β-strands of Dendra unfolded during translocation. **(E)** Cysteine-free Dendra substrate was polyubiquitinated with ubiquitin carrying a Cys at position 3, resulting in Ub^I3C^(n)-Dendra. Ub^I3C^(n)-Dendra was then incubated with different combinations of Npl4, Ufd1, and full-length Cdc48 (Cdc48^FL^) or Cdc48 containing only the N and D1 domains (Cdc48^ND1^). The samples were incubated with a maleimide-conjugated fluorescent dye (Dy-maleimide) and analyzed by SDS-PAGE, followed by fluorescence scanning (upper panel) and Coomassie-blue staining (lower panel). Ubiquitin modification was quantitated by measuring fluorescence intensities (numbers under the lanes). **(F)** Ub(n)-Dendra carrying a fluorescent dye was incubated in the presence of ADP with different combinations of Ufd1, Npl4-FLAG, and Cdc48. FLAG antibody beads (Anti-FLAG) were added, and bound material analyzed by SDS-PAGE, followed by fluorescence scanning (upper panel) and Coomassie-blue staining (lower panel). To evaluate the pull-down efficiency, different amounts of the input material were loaded (left four lanes).

We first performed HDX experiments after incubating photoconverted Ub(n)-Dendra with Cdc48 complex in the presence of ADP, i.e. conditions in which Dendra is not translocated by the ATPase and remains folded, while the initiator ubiquitin should be unfolded. Deuterium was then added and HDX was followed for different time periods (**Figure 2B**). The reaction was quenched by low pH, and the proteins proteolytically cleaved. Peptides derived from ubiquitin and Dendra were analyzed by MS and the extent of deuterium labeling determined. Analysis of ubiquitin peptides showed very little Cdc48 complex-dependent labeling at early time points of HDX (**Figure 2C**). With time, however, significantly more HDX was observed in the presence of Cdc48 complex, consistent with the complex causing ubiquitin unfolding. Deuteration was particularly strong in an N-terminal ubiquitin segment and a region in the middle of the sequence, which in the cryo-EM structure interact with Cdc48 and the Npl4 groove, respectively (Twomey et al., 2019). Ultimately, almost the entire ubiquitin population became labeled (**Figure S2A**), suggesting that most ubiquitin molecules in a polyubiquitin chain can serve as initiators and undergo Cdc48 complex-induced cycles of unfolding and refolding. Dendra peptides showed little labeling in the presence of ADP (**Figure 2D**). In contrast, when the preincubation was performed in the presence of ATP, so that substrate was unfolded before HDX (**Figure 2B**), two N-terminal β-strands of Dendra were maximally deuterated even at the shortest labeling time (**Figure 2D**). This is consistent with the expectation that, during the preincubation, the N-terminal fragment of Dendra (residues 1-192) is translocated through the central pore, while the C-terminal fragment (residues 193-355) stays behind, thus separating the two β-strands and allowing their deuteration. Surprisingly, the presence of ATP during the unfolding reaction had no effect on the subsequent deuteration of ubiquitin peptides (**Figure 2C**). As shown below, ubiquitin molecules are actually translocated through the central pore and transiently unfolded, but they refold and again serve as initiators (**Figure 7A**).

To directly measure ubiquitin unfolding, we mutated Ile3 of ubiquitin to Cys (Ub^I3C^). According to the crystal structure of ubiquitin (PDB code 1UBQ), the side chain of this residue is buried in the interior of the folded molecule and should only be exposed after unfolding. Unfolding was examined in the presence of ADP by modification of Ub^I3C^ with a maleimide-conjugated fluorescent dye (Dy-maleimide), employing Ub(n)-Dendra substrate that lacks other cysteines (**Figure 2E**). Some weak modification occurred even in the absence of Cdc48 or its cofactors (**Figure 2E**; lane 1), suggesting occasional spontaneous unfolding of ubiquitin. Ufd1 or Npl4 alone had little effect (lanes 2 and 3), but together moderately stimulated modification (lane 5). Modification was strongest when Cdc48 was also present (lane 8). Thus, all components of the Cdc48 complex are required for efficient ubiquitin unfolding.

Cooperation of the Cdc48 components was also seen in binding experiments (**Figure 2F**). For these experiments, we employed a fusion of the N-end-rule degron with super-folder GFP (sfGFP) and attached a fluorescent dye to a Cys in the N-terminal segment for better detection (Ub(n)-Dy-sfGFP) (Bodnar and Rapoport, 2017b). The Cdc48 complex was captured through FLAG-tagged Npl4 and FLAG-antibody beads, and bound proteins analyzed by SDS-PAGE (**Figure 2F**). Binding of Ub(n)-Dy-sfGFP was strongest when all components were present (lane 12). Much less binding was seen in the absence of Cdc48 or Ufd1 (lanes 10 and 11). Quantification of the pull-down efficiencies indicated that about 50% of all fully assembled Cdc48 complexes contain bound Ub(n)-Dy-sfGFP. A stimulatory effect of Cdc48 on substrate binding was also seen when a streptavidin-binding peptide (SBP) tag was attached to Ufd1 and the pull-down experiments were performed with streptavidin beads (**Figure S2B**). These results show that Npl4 and Ufd1 are not sufficient for optimal binding of polyubiquitinated substrates. Previous experiments suggested that they contain all known ubiquitin binding activity (Park et al., 2005; Sato et al., 2019), but our experiments reveal that when the cofactor components are present at equimolar concentrations, the Cdc48 ATPase greatly contributes to substrate binding.

### Insertion of the N-terminal segment of the initiator ubiquitin into the Cdc48 ring

The cryo-EM structure raised the possibility that Cdc48 stimulates substrate binding by capturing the N-terminal segment of the initiator ubiquitin in the central pore of the D1 ring. Indeed, a truncated Cdc48 protein containing only the N and D1 domains (Cdc48^ND1^) stimulated ubiquitin unfolding in the Ub^I3C^-labeling assay (**Figure 2E**, lanes 11-20) and the binding of polyubiquitinated substrate to the ATPase complex (**Figure S2C**). Capture of the unfolded ubiquitin in the D1 pore is further supported by site-specific crosslinking experiments, in which the photoreactive probe p-benzoyl-phenylalanine (Bpa) was incorporated by amber-codon suppression (Chin et al., 2002) into the D1 ring of either full-length Cdc48 or Cdc48^ND1^; in both cases, crosslinks to polyubiquitinated substrate were observed in ADP or ATP (**Figure S2D**). Taken together, these results show that the D2 ATPase ring is not required for ubiquitin unfolding and binding.

Next, we tested whether the insertion of the initiator ubiquitin segment requires the pore loops of the D1 ring. Binding experiments with Ub(n)-Dy-Dendra showed that a Cdc48^ND1^ mutant lacking the pore loop residues (ΔD1Loops) had a reduced affinity compared to wild-type Cdc48^ND1^ or a mutant of Cdc48^ND1^ that lacks ATPase activity (E315A) (**Figure 3A**; lane 5 versus lanes 4 and 6). The importance of the D1 pore loops is further highlighted by substrate unfolding experiments that utilized photoconverted Ub(n)-Dy-Dendra; in this assay, the two fragments of photoconverted Dendra are separated during the unfolding reaction, which results in a loss of fluorescence. Almost complete substrate unfolding was observed with wild-type Cdc48 complex or the E315A mutant (**Figure 3B**), consistent with D1 ATPase activity not being required (Blythe et al., 2017; Bodnar and Rapoport, 2017b). However, mutant Cdc48 lacking the D1 pore loops was completely inactive (**Figure 3B**), highlighting the importance of an interaction between the D1 pore loops and the N-terminal segment of the initiator ubiquitin molecule.

**Figure 3.**
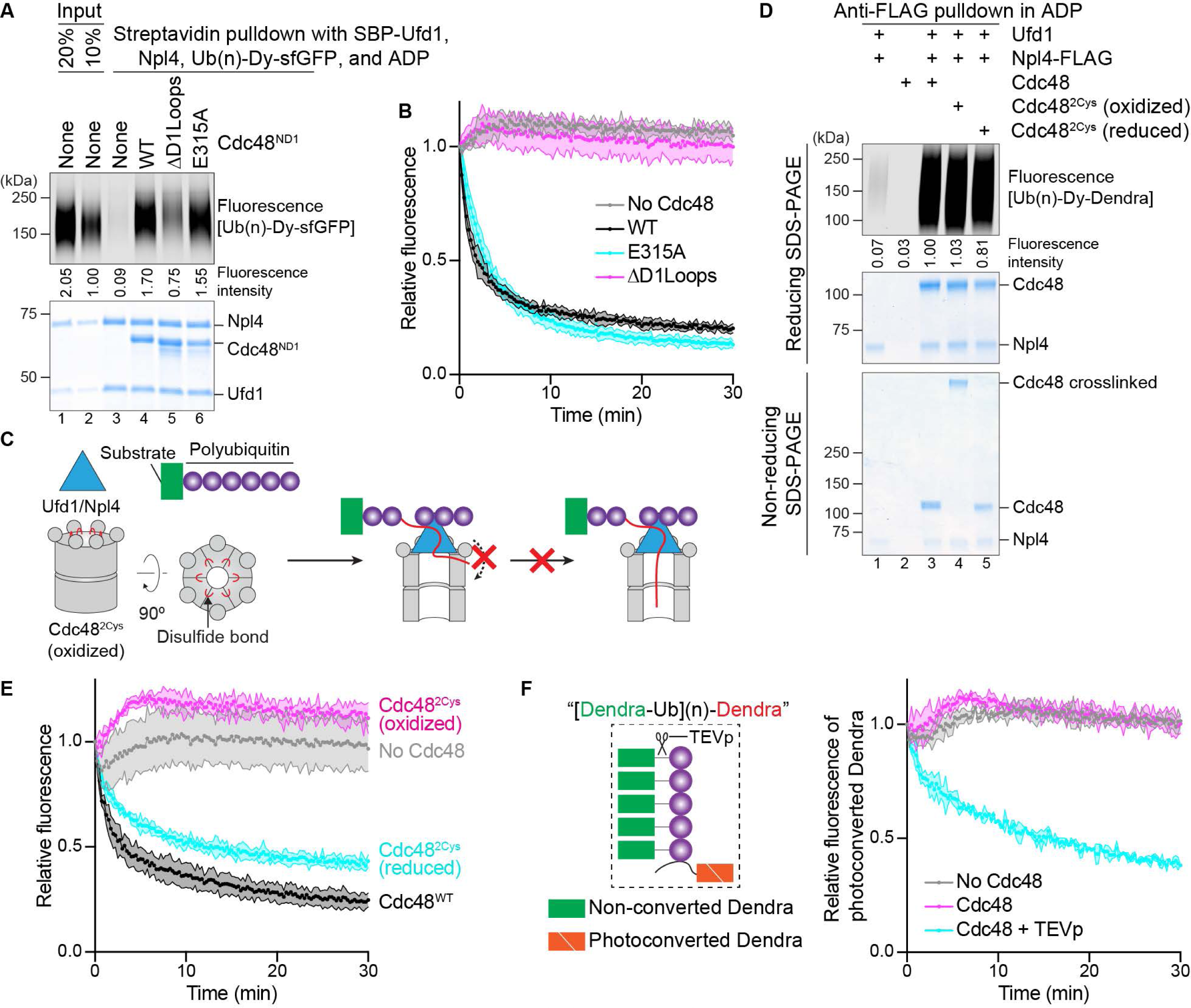
Insertion of the N-terminal segment of the initiator ubiquitin into the D1 ATPase ring. **(A)** Cdc48 lacking the D2 domain (Cdc48^ND1^), which was otherwise wild-type (WT), lacked the pore loops in the D1 domain (ΔD1loops), or was deficient in ATPase activity (E315A), was incubated with SBP-tagged Ufd1 (SBP-Ufd1), Npl4, and dye-labeled Ub(n)-sfGFP in the presence of ADP. The Cdc48 complex was retrieved with streptavidin beads, and bound material analyzed by SDS-PAGE, followed by fluorescence scanning (upper panel) and Coomassie-blue staining (lower panel). Substrate was quantitated by measuring fluorescence intensities (numbers under the lanes). **(B)** Photoconverted Ub(n)-Dendra was incubated in the presence of ATP with Cdc48/UN complex containing full-length wild-type (WT) Cdc48, or the D1 mutants ΔD1loops or E315A. Unfolding of Dendra was followed by the loss of fluorescence. The experiments were performed in triplicates. Shown are means and standard deviations for each data point. **(C)** Scheme for testing whether the N-terminus of the initiator ubiquitin enters the D1 ring from the side. Lateral entry is prevented by crosslinking all six D1 domains in the hexameric ring through disulfide bridges positioned close to the entrance of the pore. **(D)** Crosslinked Cdc48 hexamers were generated with Cdc48 containing S286C and R333C (Cdc48^2Cys^) by addition of an oxidant (oxidized). One aliquot was treated with a reducing reagent to generate non-crosslinked hexamers (reduced). The samples were incubated with Ufd1, Npl4-FLAG, and photoconverted dye-labeled Ub(n)-Dendra in the presence of ADP, followed by retrieval of the Cdc48 complex with FLAG-antibody beads. As a control, wild-type Cdc48 was used. Bound material was analyzed by reducing and non-reducing SDS-PAGE, followed by fluorescence scanning and Coomassie-blue staining. **(E)** Cdc48/UN complex containing wild-type Cdc48 (Cdc48^WT^), Cdc48^2Cys^ (oxidized), or Cdc48^2Cys^ (reduced) was tested for unfolding of photoconverted Ub(n)-Dendra, as in (B). **(F)** The ubiquitin chain in photoconverted Ub(n)-Dendra was replaced with a chain of Dendra-ubiquitin fusions containing a TEV protease cleavage site between the fusion partners ([Dendra-Ub](n)-Dendra; see scheme). Note that Dendra in the fusions was not photoconverted. After incubation with or without TEV protease (TEVp), unfolding of photoconverted Dendra was tested in the absence or presence of Cdc48 complex, as in (B).

Given the substantial length of the N-terminal segment of the initiator ubiquitin and its extended conformation in the cryo-EM structure, it seemed possible that it would enter the central pore of the D1 ring from the side. To test this possibility, we generated a Cdc48 hexamer in which all subunits are disulfide-crosslinked to their neighbors so that lateral entry would be prevented (**Figure 3C**). This was achieved by introducing cysteines at S286 and R333, which are located at the interface of neighboring protomers close to the D1 pore entrance (**Figure S2E**). Treatment of this mutant (Cdc48^2Cys^) with an oxidant resulted in efficient crosslinking of all six protomers of the Cdc48 hexamer (**Figure 3D**; bottom panel). The crosslinked Cdc48^2Cys^ complex stimulated binding of polyubiquitinated substrate to the UN cofactor to a similar extent as the non-crosslinked complex (**Figure 3D**; upper most panel; lane 4 versus lanes 3 and 5). Thus, the N-terminal segment of the initiator ubiquitin likely enters the pore from the top of a closed D1 ring, rather than from the side. Interestingly, the crosslinked Cdc48^2Cys^ hexamer was inactive in substrate unfolding, but was re-activated upon reduction of the disulfides (**Figure 3E**). Given that ATP hydrolysis in the D1 protomers is not required for substrate unfolding (**Figure 3B**; E315A mutant), we consider it unlikely that crosslinking simply prevented movements of the D1 protomers; rather, a closed hexameric D1 ring may prevent the K48-branchpoint of the initiator ubiquitin from moving through the central pore (see below).

To further test whether the N-terminus of the initiator ubiquitin enters a closed D1 ring, we generated a substrate in which the ubiquitin chain in photoconverted Ub(n)-Dendra was replaced with a chain of Dendra-ubiquitin fusions (**Figure 3F**); the bulky Dendra domains should prevent the N-termini of all ubiquitin molecules from entering the ATPase ring. Indeed, no unfolding was observed with this substrate (**Figure 3F**). However, when Dendra was removed by taking advantage of a TEV protease cleavage site between the fusion partners, unfolding was restored (**Figure 3F**). These experiments show that a free N-terminus is required for ubiquitin to serve as initiator. Interestingly, when the small ubiquitin-like modifier (SUMO) protein was used as N-terminal fusion partner of ubiquitin, slow unfolding was observed (which was accelerated after cleavage of SUMO by the Ulp1 protease) (**Figure S2F**). A likely explanation is that a SUMO molecule can serve as initiator, albeit not as efficiently as ubiquitin.

Taken together, our results show that the Cdc48 ATPase has a hitherto unappreciated role in substrate recruitment: it uses the pore loops of the D1 ring to capture the N-terminal segment of the unfolded initiator ubiquitin inside the central pore, thereby augmenting the binding of ubiquitin molecules to the Cdc48/UN complex.

### Ubiquitin unfolding requires the cofactors Npl4 and Ufd1

Next, we tested the role of the cofactors in ubiquitin unfolding. We first made several mutations in the Npl4 groove (**Figure S3A**), which according to the cryo-EM structure (Twomey et al., 2019), accommodates residues 23 to 48 of the initiator ubiquitin. Mutation of amino acids at the bottom or top of Npl4’s groove (**Figure S3A**; inset i) reduced the binding of Ub(n)-Dy-Dendra to the Cdc48 complex, whereas mutation of residues contacting the kink of the initiator ubiquitin (**Figure S3A**) had little effect (**Figure S3B**). Binding of Ub(n)-Dy-Dendra to the mutants showed a correlation with the initial rate of Dendra unfolding (**Figure S3C**). Ub(n)-Dy-Dendra binding also correlated with the efficiency of polypeptide insertion into the central pore of Cdc48, as demonstrated by site-specific photo-crosslinking experiments with Bpa probes in the D2 ring of Cdc48 (**Figure S3D**). These results indicate that the Npl4 groove plays an important role in the binding of the initiator ubiquitin and subsequent substrate processing.

We next probed the interaction of Npl4 with the two folded ubiquitin molecules Ub1 and Ub2 visible in the cryo-EM structure (**Figure S3A**; insets ii and iii). Mutations designed to disrupt the interaction with Ub2 (Sato et al., 2019) only moderately reduced Ub(n)-Dy-sfGFP binding (**Figure S3E**; lanes 5-7 versus 4). Two of the four mutations introduced at the interface to Ub1 also attenuated binding (lanes 8-11). However, surprisingly, all mutations only slightly reduced substrate unfolding (**Figure S3F**). Thus, the interaction of Npl4 with the folded ubiquitin molecules is not crucial, consistent with the low sequence conservation of Npl4 in the interacting regions (Twomey et al., 2019). These results suggest that Ufd1 may be more important for the interaction with folded ubiquitin molecules.

To test the role of Ufd1, we deleted the N-terminal, ubiquitin-binding UT3 domain (Park et al., 2005) (**Figure S3G**). Indeed, substrate recruitment to the Cdc48 complex was completely abolished (**Figure S3H;** lane 4 versus 5). The same result was obtained when photocrosslinking of polyubiquitinated substrate to Cdc48’s D2 ring was tested (**Figure S3I;** lane 4 versus 3), or when the unfolding of photoconverted Dendra was measured (**Figure S3J**). Thus, the ubiquitin binding activity of the UT3 domain of Ufd1 plays an essential role in substrate recruitment. Taken together, these results show that the ubiquitin binding activities of the UN cofactor promote the unfolding of the initiator ubiquitin and substrate recruitment to the Cdc48 ATPase.

### Translocation of the initiator ubiquitin

Next, we investigated the translocation phase. We first tested whether translocation begins with the Cdc48 ATPase pulling on the initiator ubiquitin. In this case, the initiator segment originally bound to the Npl4 groove should be dislodged, which in turn should lead to the displacement of Ub1 and Ub2 from the top of the Npl4 tower; if these ubiquitin molecules cannot be released from their binding sites, translocation and substrate unfolding should be prevented (**Figure S4A**). To test this prediction, we incorporated photoreactive Bpa probes into the Npl4 groove and the top of the Npl4 tower by amber-codon suppression (**Figure 4A**). The Bpa-containing Npl4 mutants were mixed with photoconverted Ub(n)-Dendra, Ufd1, and Cdc48, and the mixtures were irradiated with UV light in the absence of nucleotides; ATP was then added and the unfolding of Dendra monitored. All tested positions could crosslink to polyubiquitinated substrate (**Figure S4B**), and all crosslinked complexes showed a reduced ability to unfold substrate (**Figure 4B**). In contrast, irradiated mixtures containing wild-type Npl4 and all non-irradiated samples showed unperturbed unfolding kinetics (**Figure 4B**). In general, there was a good correlation between the crosslinking yield and inhibition of the initial unfolding rate (**Figure 4C**), indicating that the crosslinked complexes were entirely inactive. These results show that translocation is prevented when the ubiquitin molecules cannot be released from Npl4, consistent with translocation beginning with the N-terminal segment of the initiator ubiquitin and causing the initiator ubiquitin, Ub1, and Ub2 to be dislodged from their original binding sites on Npl4.

**Figure 4.**
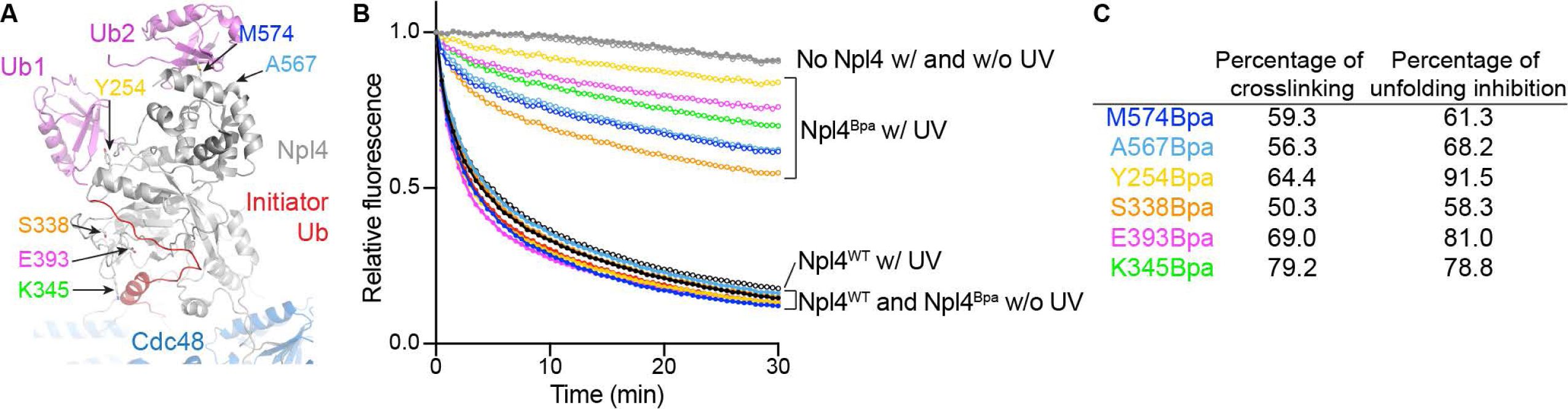
Translocation of the initiator ubiquitin by Cdc48. **(A)** Model of Npl4 with bound ubiquitin molecules, based on a cryo-EM structure of Cdc48 in complex with polyubiquitinated substrate (PDB, 6OA9). Npl4 is shown in grey, the unfolded initiator ubiquitin molecule in red, the two folded ubiquitin molecules Ub1 and Ub2 in pink, and Cdc48 in blue. Bpa probes were incorporated at the indicated Npl4 positions. **(B)** The unfolding of photoconverted Ub(n)-Dendra was tested with Cdc48 complex containing wild-type Npl4 (Npl4^WT^) or Npl4 with Bpa probes at positions interacting with the initiator ubiquitin, Ub1, or Ub2 (Npl4^Bpa^; the curve colors correspond to the positions shown in (A)) with (w) or without (w/o) UV-induced crosslinking. **(C)** For each Bpa mutant, the percentage of substrate crosslinked to Npl4 was determined (**Figure S4B**) and compared with the percentage of inhibition of the initial rate of Dendra unfolding.

### Translocation of polypeptide branchpoints by the Cdc48 ATPase

The next important event happens when the K48 branchpoint of the initiator ubiquitin (called type I) enters the ATPase rings (**Figure 5A**), as the ATPase could in principle translocate either polypeptide branch through its central pore. Our unfolding experiments with Ub(n)-Dendra (e.g. **Figure 3B**) indicate that Cdc48 can move towards the C-terminus of the initiator ubiquitin, as this is the direction in which it can reach the substrate that is either directly attached to the C-terminus or separated from the initiator by proximal ubiquitin molecules (**Figure 5A**).

**Figure 5.**
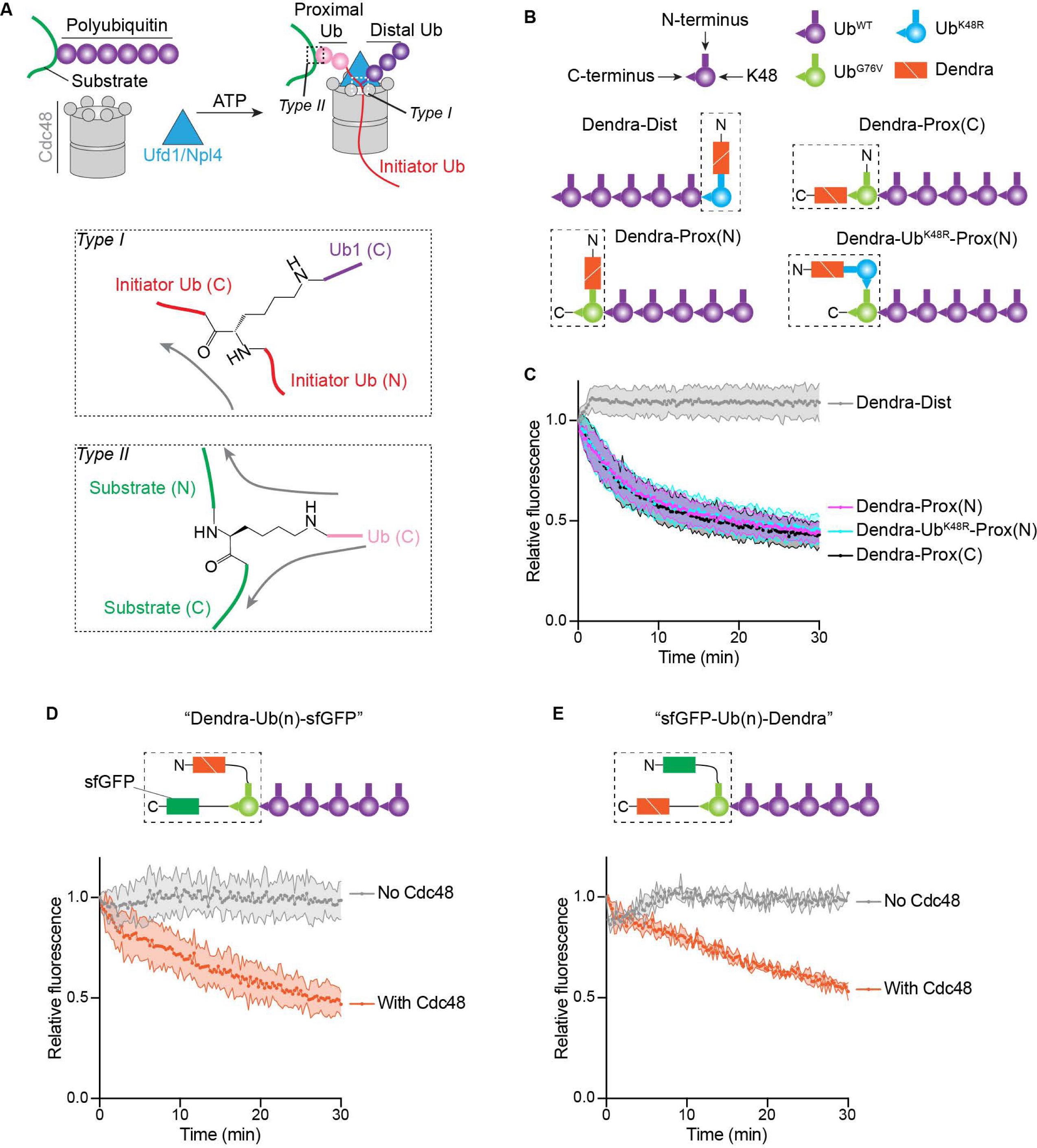
Translocation of branched polypeptides by the Cdc48 ATPase. **(A)** Scheme of substrate processing by the Cdc48 ATPase complex. Ubiquitin molecules proximal and distal from the initiator (in red) are indicated in pink and purple, respectively. The two different branchpoints (type I and II) are indicated by dashed boxes in the upper scheme and magnified in the lower panels. Arrows indicate the directions of translocation through the Cdc48 pore. **(B)** Schemes of different polyubiquitinated substrates used for unfolding experiments. Proteins linked by fusion are shown in a dashed box, with N- and C-terminus indicated. Dist and Prox indicate that photoconverted Dendra is located either at the distal or proximal end of a polyubiquitin chain. (N) and (C) indicate that Dendra is located at the N- or C-terminus of the fused ubiquitin. **(C)** The substrates shown in (B) were tested for unfolding of photoconverted Dendra by measuring the loss of fluorescence. The experiments were performed in triplicates. Shown are means and standard deviations for each data point. **(D)** A polyubiquitinated substrate was generated with a fusion protein containing photoconverted Dendra, ubiquitin, and sfGFP (dashed box) and tested for the unfolding of Dendra. **(E)** As in (D), but with swapped positions of sfGFP and Dendra in the fusion protein.

To test whether Cdc48 can also translocate in the other direction when it encounters the K48 branchpoint of the initiator and therefore unfold distal ubiquitins, we generated a K48-linked polyubiquitin chain with photoconverted Dendra attached to the most distal ubiquitin molecule (**Figure 5B**; Dendra-Dist). This was achieved by performing a ubiquitination reaction with a mixture of wild-type ubiquitin (Ub^WT^) and a fusion of Dendra to the K48R mutant of ubiquitin (Dendra-Ub^K48R^), using a purified fusion enzyme consisting of the RING finger domain of the ubiquitin ligase gp78 and the conjugating enzyme Ube2G2 (Blythe et al., 2017); incorporation of Ub^K48R^ terminates chain elongation, resulting in Dendra-Ub^K48R^ capping the ubiquitin chains. Dendra-Dist was able to bind to the Cdc48 complex (**Figure S4C**), but addition of ATP did not cause any reduction of Dendra fluorescence (**Figure 5C**), indicating that Cdc48 does not unfold distal ubiquitin molecules. As a control, we generated a similar substrate, but with photoconverted Dendra at the proximal side of the initiator ubiquitin (**Figure 5B**; Dendra-Prox(C)). Like Ub(n)-Dendra (**Figure 3B**), Dendra-Prox(C) substrate was efficiently unfolded (**Figure 5C**), again showing that proximal ubiquitins are translocated through the central pore of Cdc48.

The next crucial event happens when the ATPase reaches the C-terminus of the initiator, as it encounters another branchpoint (called type II), either K48 of another ubiquitin molecule or a Lys residue of the substrate. This branchpoint is fundamentally different from the K48 branchpoint of the initiator (type I), as Cdc48 is now pulling on the isopeptide bond of a folded protein, rather than on the peptide bond of an unfolded ubiquitin segment protein (**Figure 5A**). Again, we asked whether Cdc48 translocates one or both branches through its central pore. Because photoconverted Dendra was attached to the C-terminus of the most proximal ubiquitin molecule in Ub(n)-Dendra, its unfolding indicates that the ATPase can translocate from the type II branchpoint towards the C-terminus of the substrate. To test whether Cdc48 can also translocate in the other direction, we fused photoconverted Dendra to the N-terminus of the ubiquitin mutant Ub^G76V^ and attached a K48-linked ubiquitin chain to Ub^G76V^ (**Figure 5B**; Dendra-Prox(N)); the G76V mutation ensures that the fusion protein is at the proximal end of the ubiquitin chain and that the K48 of Ub^G76V^ is a type II branchpoint. Dendra was efficiently unfolded by Cdc48 (**Figure 5C**), demonstrating that, in contrast to the type I branchpoint, type II allows bidirectional translocation towards either the N- or C-terminus. One possible explanation for the selectivity at type I branchpoints is that Cdc48 cannot unfold ubiquitin when it pulls on its C-terminus. To test this possibility, we generated a substrate that contained Dendra fused to Ub^K48R^ at the N-terminus of a polyubiquitin chain (**Figure 5B**; Dendra-Ub^K48R^-Prox(N)). In this case, translocation can only be initiated within the ubiquitin chain, and when Cdc48 reaches Ub^K48R^ (blue), it has to pull on the C-terminus of this ubiquitin molecule before it can unfold Dendra. Nevertheless, Dendra was efficiently unfolded (**Figure 5C**; Dendra-Ub^K48R^-Prox(N)). Thus, Cdc48 can unfold ubiquitin when it pulls on its C-terminus, but not at type I branchpoints.

To further test whether Cdc48 can translocate in either direction when it encounters a type II branchpoint, we generated fusion proteins in which ubiquitin is flanked by sfGFP and photoconverted Dendra (Dendra-Ub(n)-sfGFP and sfGFP-Ub(n)-Dendra) (**Figures 5D; 5E)**. Both substrates were unfolded by the Cdc48 complex. Translocation of these substrates cannot be initiated by the ubiquitin molecule in the fusion protein because its N-terminus is blocked by a bulky protein (see **Figure 3F**). Rather, one of the ubiquitin molecules in the attached chain (shown in purple) must have served as initiator, and translocation must have proceeded through the ubiquitin in the fusion protein to Dendra. These results confirm that Cdc48 can translocate towards either the N- or C-terminus when it encounters the isopeptide bond of the substrate-attached ubiquitin molecule. Taken together, our results indicate that at the type I branchpoint of the initiator ubiquitin, Cdc48 selectively translocates the C-terminal segment through the central pore, but at subsequent type II branchpoints, it can move either polypeptide branch through the pore.

### Release of polyubiquitinated substrate from Cdc48

Finally, we investigated whether the polyubiquitinated substrate is released from the Cdc48 complex after its translocation through the central pore. Ub(n)-Dendra was associated with the Cdc48 complex, regardless of whether the incubation was performed in ADP or ATP (**Figure 6A**, lane 5 versus 6), i.e. whether or not Dendra was folded or unfolded (**Figure S5A**). Thus, fully translocated substrate was either not released from the Cdc48 complex or re-associated with it (**Figure 6B**; scenarios a and b, respectively). Substrate release would be prevented if the distal ubiquitins stayed on the cis side of the ATPase ring, such that proximal and distal ubiquitins would end up on opposite sides of the ATPase rings, with a connecting polypeptide segment spanning the pore (**Figure 6B**; scenario a). On the other hand, if the distal ubiquitins moved outside the pore, while the K48 branchpoint of the initiator itself moved through the pore, at the end of translocation all ubiquitin molecules and substrate would be released from the Cdc48 complex and could re-associate to re-generate an initiation state, but this time with unfolded substrate (**Figure 6B**; scenario b).

**Figure 6.**
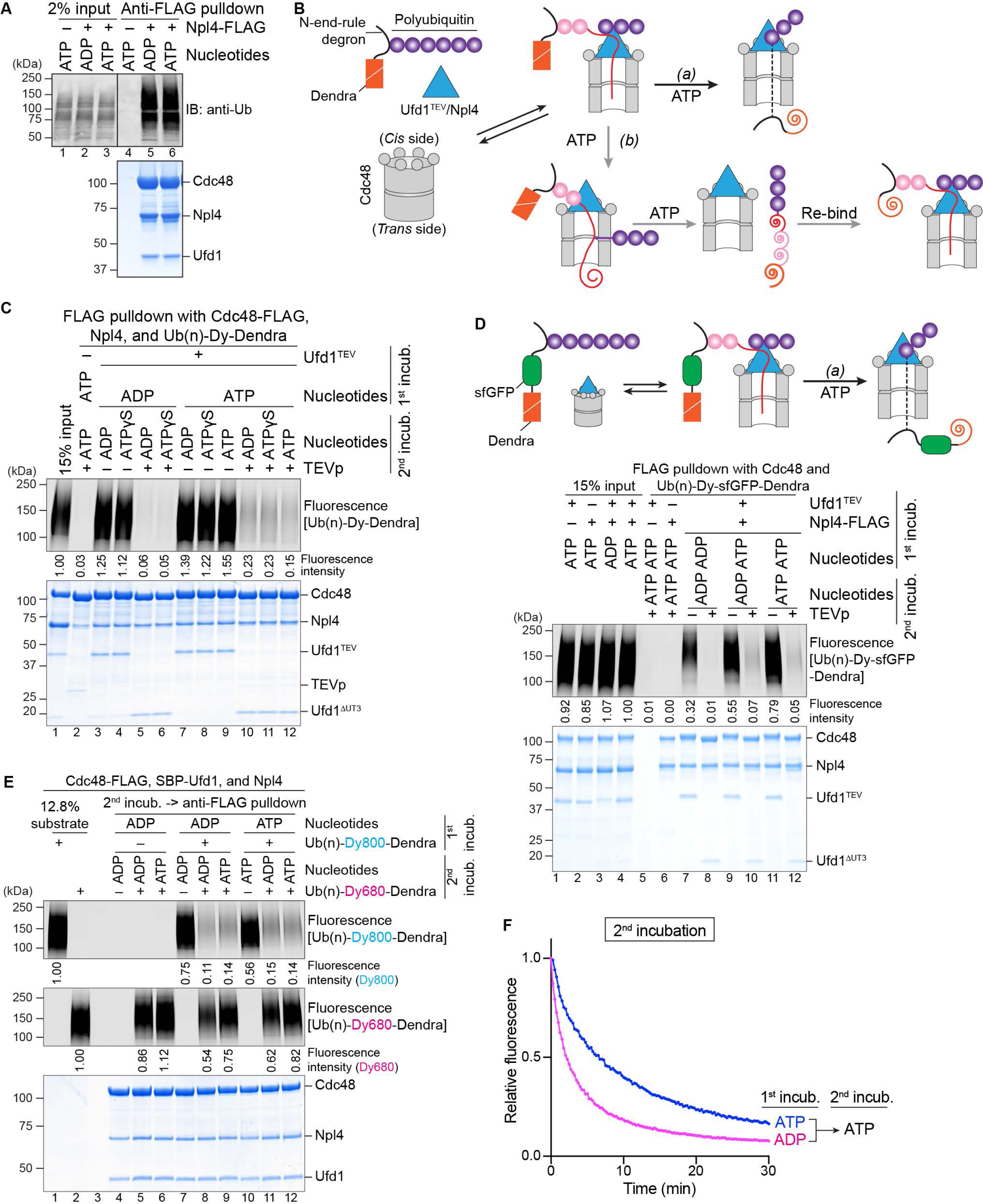
Release of polyubiquitinated substrate from the Cdc48 ATPase complex. **(A)** Photoconverted Ub(n)-Dendra, Cdc48, and Ufd1 were incubated with or without FLAG-tagged Npl4 in the presence of ADP or ATP. The Cdc48 complex was isolated with FLAG-antibody beads, and bead-bound material analyzed by SDS-PAGE, followed by immunoblotting (IB) with ubiquitin antibodies (anti-Ub; upper panel) and Coomassie-blue staining (lower panel). **(B)** Scheme showing two conceivable scenarios for the post-translocation state. In scenario (a), the distal ubiquitins (in purple) remain on the *cis* side of the ATPase ring, resulting in a polypeptide segment spanning the pore (dashed line). In this case, the substrate cannot be released by removal of the UN cofactor. In scenario (b), distal ubiquitin molecules move outside the ATPase rings, while the K48 branchpoint of the initiator ubiquitin (in red) moves through the pore. At the end, the unfolded, polyubiquitinated substrate is released from Cdc48 and can rebind to the ATPase complex. **(C)** Dye-labeled Ub(n)-Dendra was incubated in the presence of ADP or ATP with Cdc48, Npl4-FLAG, and Ufd1 containing a TEV cleavage site following the UT3 domain (Ufd1^TEV^) (1^st^ incubation). The Cdc48 complex was retrieved with FLAG-antibody beads, and bound material was treated with TEV protease in the presence of ADP, ATP, or ATPγS, a non-hydrolyzable ATP analog (2^nd^ incubation). Bead-bound material was then analyzed by SDS-PAGE, followed by fluorescence scanning (upper panel) and Coomassie-blue staining (lower panel). **(D)** As in (C), but with Ub(n)-sfGFP-Dendra. Note that sfGFP refolds after translocation, which would prevent backsliding of the polypeptide through the Cdc48 pore in scenario (a) (upper panel). **(E)** Photoconverted Ub(n)-Dendra labeled with the fluorophore DyLight800 (Ub(n)-Dy800-Dendra) was incubated with Cdc48-FLAG, SBP-Ufd1, and Npl4 in the presence of ADP or ATP (1^st^ incubation). The Cdc48 complex was retrieved with streptavidin beads and eluted with biotin. The samples were then incubated with DyLight680-labeled, photoconverted Ub(n)-Dendra (Ub(n)-Dy680-Dendra) in the presence of ADP or ATP. Anti-FLAG antibody beads were added, and bound material analyzed by SDS-PAGE, followed by fluorescence scanning at 800nm and 680nm wavelength (upper two panels) and Coomassie-blue staining (lower panel). **(F)** Samples in (E), incubated in ADP or ATP during the first incubation, were tested for unfolding of photoconverted Ub(n)-Dy680-Dendra in ATP during the second incubation (Figure 6E; lanes 9 versus 12).

**Figure 7.**
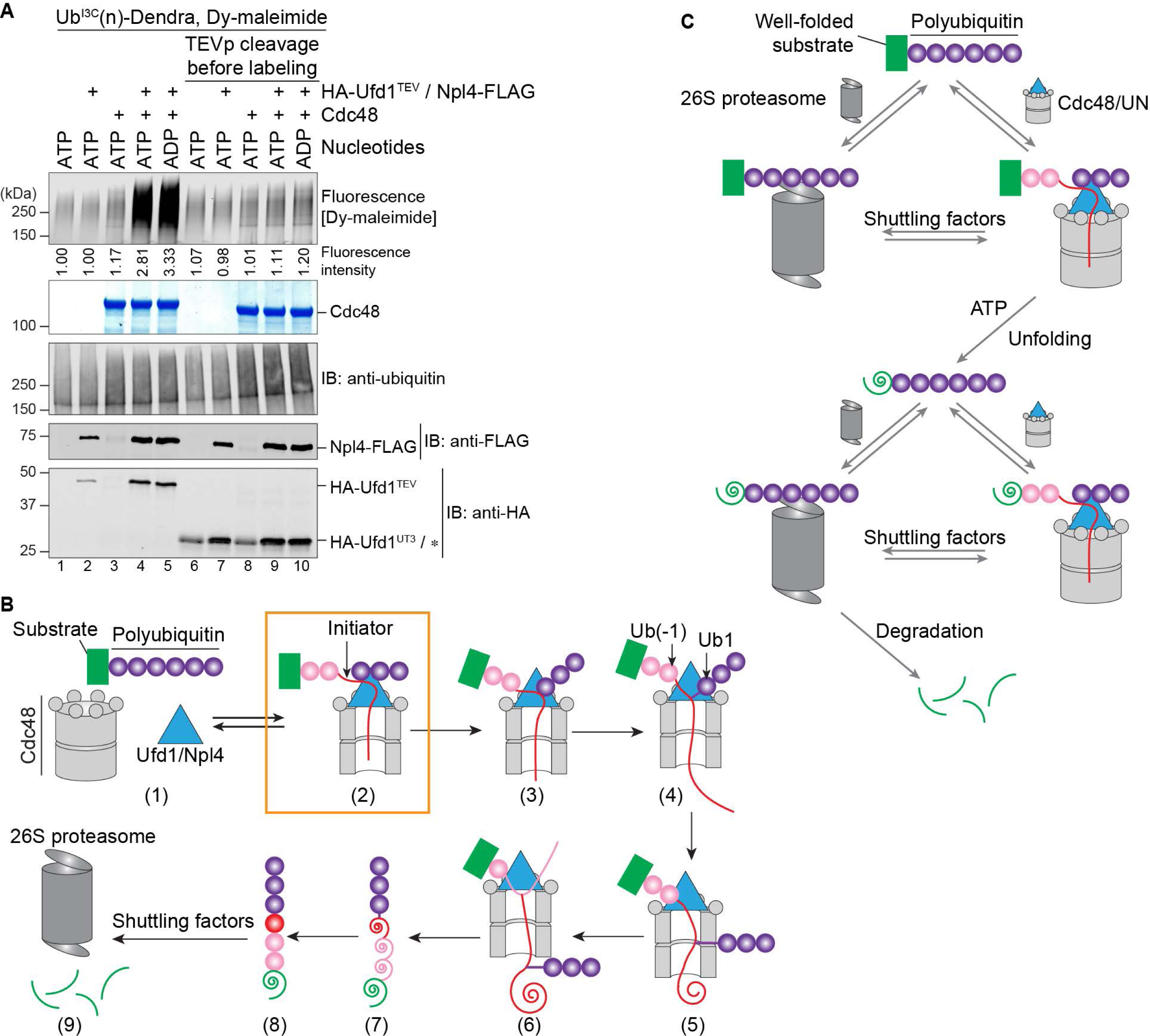
Refolding of translocated ubiquitin and model for substrate processing by the Cdc48 complex. **(A)** Ub^I3C^(n)-Dendra was incubated in the presence of ADP or ATP with Cdc48-FLAG, Npl4-FLAG, and HA-tagged Ufd1 containing a TEV protease cleavage site following the UT3 domain (HA-Ufd1^TEV^). Where indicated, TEV protease (TEVp) was added after incubation with the nucleotides. All samples were then incubated with a maleimide-conjugated fluorescent dye (Dy-maleimide) and analyzed by SDS-PAGE, followed by fluorescence scanning (upper panel), Coomassie-blue staining (2^nd^ panel), and immunoblotting (IB) with ubiquitin, FLAG, and HA antibodies. The HA antibodies cross-react with TEV protease, which migrates at the same position as UT3 (star). **(B)** Scheme of substrate processing by the Cdc48 ATPase complex. The boxed initiation state is most populated. The unfolded initiator ubiquitin is shown as a red line, and proximal and distal ubiquitin molecules as pink and purple circles, respectively. Translocated, unfolded ubiquitin and substrate molecules are indicated as spirals. Unfolded substrate released from the Cdc48 complex can be transferred by shuttling factors to the 26S proteasome and be degraded. **(C)** Model for the transfer of polyubiquitinated proteins between the Cdc48 complex and the 26S proteasome. Note that both particles and the shuttling factors all recognize primarily the ubiquitin chain.

To test whether the final state resembles an initiation state, we took advantage of the fact that polyubiquitinated substrate cannot bind to the Cdc48 complex without the UT3 domain of Ufd1 (**Figure S3H**). We generated a Cdc48 complex, in which Ufd1 contains a TEV cleavage site following its UT3 domain (Ufd1^TEV^), a complex that was active in Dendra unfolding (**Figure S5B**). Ub(n)-Dendra, bound to this Cdc48 complex in ADP (first incubation), was released after incubation with TEV protease (**Figure 6C**; lanes 5 and 6 versus lanes 3 and 4), confirming that the UT3 domain is required for the interaction. When the first incubation was performed in ATP to induce Dendra unfolding (**Figure S5B**), removal of the UT3 domain also caused Ub(n)-Dendra to dissociate from the Cdc48 complex (**Figure 6C**; lane 10-12 versus 7-9), consistent with the idea that unfolded Ub(n)-Dendra was released from the Cdc48 complex after translocation and re-associated with it as in the initiation state (scenario b).

To further exclude scenario a, we used a polyubiquitinated fusion protein containing both sfGFP and photoconverted Dendra ((Ub(n)-sfGFP-Dendra) (**Figure 6D**). Photoconvered Dendra, consisting of two non-covalently associated fragments, is irreversibly unfolded by Cdc48 (**Figure S5C**), while the single-polypeptide domain of sfGFP can rapidly refold after its translocation (**Figure S5D**). If a polypeptide segment spanned the pore (scenario a), the refolded, bulky sfGFP domain would prevent backsliding of the substrate through the central pore, and the Cdc48 pore loops would interact with the polypeptide segment located inside the central pore. In this case, polyubiquitinated substrate would remain associated with the Cdc48 complex after UT3 removal. However, the results show that the association of Ub(n)-sfGFP-Dendra with Cdc48/UN was drastically reduced whenever the UT3 domain was cleaved off (**Figure 6D**), regardless of whether Dendra was folded or unfolded (ADP or ATP in the first incubation). Because dissociation after TEV protease cleavage occurred even when the second incubation was performed in ADP (**Figure 6D**, lane 10), translocation is not required for substrate dissociation following UT3 removal.

To further test whether folded and unfolded polyubiquitinated substrates are bound to the Cdc48 complex in the same way, we performed competition experiments (**Figure 6E**). Dy800-labeled Ub(n)-Dendra was incubated with Cdc48 complex in either ADP or ATP to generate initiation and post-translocation complexes, respectively (**Figure 6E**; 1^st^ incubation). As expected, unfolding was only observed in ATP (**Figure S5E**). Complexes of Dy800-labeled substrate and Cdc48 complex were then isolated and incubated with an excess of Dy680-labeled Ub(n)-Dendra, again in the presence of ADP or ATP (**Figure 6E**, 2^nd^ incubation). Regardless of the conditions, the second substrate competed efficiently with the first. Competition in the presence of ADP confirms that the initial binding reaction, including the unfolding of the initiator ubiquitin, is reversible. With ATP in the second incubation, the competitor substrate was also unfolded (**Figure 6F**). These results show that polyubiquitinated substrate is bound to the Cdc48 complex in the same way before and after unfolding, implying that substrate can undergo multiple rounds of translocation. Indeed, by varying the concentrations of Cdc48 and substrate, we found that each Cdc48 hexamer can unfold more than one substrate molecule (**Figure S5F**).

Our conclusions are further supported by hydrogen/deuterium exchange (HDX) MS experiments (**Figure S6A-C**). Cdc48 complex was incubated with photoconverted Ub(n)-Dendra in ADP or ATP, and the samples were subjected to HDX for different time periods and analyzed by MS. All peptides derived from Cdc48, Npl4, and Ufd1 showed only small deuteration differences between ADP and ATP (**Figure S6A-C**), consistent with the idea that all components of the Cdc48 complex are in the same state before and after substrate unfolding.

Additional evidence for this conclusion was obtained by site-specific photocrosslinking followed by MS. Polyubiquitinated substrate was incubated in the presence of ADP or ATP with Cdc48 complex containing a photoreactive Bpa probe in the D2 ring (position 602) to generate Cdc48 complex with associated folded or unfolded substrate (**Figure S7A**). The samples were then irradiated, and the crosslinked products (**Figure S7B)** analyzed by trypsin digestion, followed by nano-liquid chromatography and MS (**Figure S7C; S7D**). In the presence of ADP, the Bpa probe crosslinked to residues Met^1^ or Gln^2^ of ubiquitin (**Figure S7C**), in agreement with previous results for the initiation state (Twomey et al., 2019). Similar results were obtained in the presence of ATP when essentially all substrate molecules were unfolded (**Figure S7D**). No other crosslinked substrate or ubiquitin peptide could be detected, probably because the dwell time of the N-terminal segment of the initiator ubiquitin in the central pore is much longer than that of any other segment.

### Refolding of translocated ubiquitin molecules

Because the pore of Cdc48 ATPase is rather narrow, all ubiquitin molecules must be in an extended conformation during translocation. To test whether these molecules refold after release from Cdc48, we again used modification of Ub^I3C^ by a maleimide-conjugated fluorescent dye (see **Figure 2E**). Cysteine-lacking substrate was polyubiquitinated with Ub^I3C^ and incubated in the presence of ADP or ATP with Cdc48 complex containing Ufd1 with a TEV protease cleavage site following the UT3 domain (Ufd1^TEV^). Without TEV cleavage, strong ubiquitin modification was observed when all components of the Cdc48 complex were present (**Figure 7A**; lanes 4 and 5 versus 1-3). Importantly, no increase in modification was observed in ATP (lane 4 versus 5), although essentially the entire substrate population was unfolded (**Figure S7E**). Thus, all ubiquitin molecules positioned between the initiator and substrate, which were unfolded when they passed through the central pore, must have refolded after translocation. When polyubiquitinated substrate was released from the Cdc48 complex by TEV cleavage of Ufd1^TEV^ (see **Figure 6C**), modification of Ub^I3C^ remained at the ground level (**Figure 7A**; lanes 6-10). This result and the similar levels of modification in ADP and ATP indicate that all proximal ubiquitin molecules refold after their translocation through the pore; the polyubiquitinated unfolded substrate rebinds to the Cdc48 complex, with an initiator ubiquitin molecule in the central pore. This conclusion is consistent with our observation that ubiquitin molecules undergo HDX with the same kinetics before and after substrate unfolding (**Figure 2C**).

## DISCUSSION

Here we have determined the molecular mechanism by which the Cdc48 ATPase complex processes its polyubiquitinated substrates. Our results lead to a model (**Figure 7B**) that explains how Cdc48 and its mammalian ortholog p97/VCP cooperate with the UN cofactor to disassemble protein complexes and extract proteins from membranes.

The ATPase complex first binds a polyubiquitinated substrate, a process that is reversible and requires no ATP hydrolysis (**Figure 7B**, stage 1 to stage 2). All three components, i.e. Cdc48, Npl4, and Ufd1, cooperate in the initial interaction with the ubiquitin chain, which results in the unfolding of one of the ubiquitin molecules. Since all ubiquitin molecules undergo HDX, it seems that the choice of the initiator ubiquitin is random. However, this ubiquitin molecule needs to be succeeded in the chain by several folded ubiquitin molecules, two of which are bound to the top of the Npl4 tower and two or more to the UT3 domain of Ufd1. Our results indicate that the UT3 interaction is more important. Despite ubiquitin being a very stable protein, it is unfolded simply by binding to the Cdc48 complex. Unfolding is initiated by thermal fluctuation, which results in the separation of the N-terminus of ubiquitin from the rest of the molecule (Irbäck et al., 2005). Next, a ubiquitin segment binds to a groove of Npl4 and the N-terminus is captured by its insertion into the D1 ring of Cdc48. Together, these interactions shift the equilibrium towards the unfolded state of ubiquitin.

During the binding step, the N-terminal segment of the initiator ubiquitin engages the D2 pore loops, as visualized in our previous cryo-EM structure (Twomey et al., 2019). We now show that subsequent ATP hydrolysis causes this segment to be pulled through the central pore (**Figure 7B**, stage 2 to 3). As a consequence, the segment of the initiator ubiquitin originally bound to the Npl4 groove is dislodged, which in turn leads to the release of the folded ubiquitin molecules Ub1 and Ub2 from their original binding sites at the top of Npl4. The UT3 domain of Ufd1 probably continues to reversibly interact with the distal parts of the ubiquitin chain.

The next important event happens when the type I branchpoint of the initiator ubiquitin enters the D1 pore (**Figure 7B**, stage 4). We demonstrate that the ATPase continues translocation towards the C-terminus of the initiator ubiquitin (stage 4 to 5), rather than translocating distal ubiquitin molecules. The selectivity may be determined by the fact that the C-terminal segment of the initiator ubiquitin is already unfolded and therefore requires less energy for translocation than the folded, K48-attached Ub1 molecule. This Ub1 molecule and all ubiquitins distal to it remain folded and outside the central pore. Because the K48 branchpoint of the initiator ubiquitin is translocated through the pore, distal ubiquitins likely move on the side of the double-ring ATPase (**Figure 7B**, stages 4-6), with a segment between K48 and the C-terminus of the adjacent ubiquitin molecule passing through lateral openings between two protomers of the hexamers. This movement may require some of the distal ubiquitin molecules to dissociate from the UN complex. Lateral opening of the two hexameric ATPase rings seems possible, as both the D1 and D2 protomers can form open spirals, in which one protomer is invisible in cryo-EM structures (Twomey et al., 2019). A requirement for lateral ring opening is supported by our observation that a disulfide-crosslinked Cdc48 hexamer can bind, but not translocate, a polyubiquitinated substrate.

After translocating the initiator ubiquitin, Cdc48 encounters the next branchpoint (**Figure 7B**, stage 6), as the C-terminus of the initiator is linked to either a lysine residue of the substrate or K48 of another ubiquitin molecule. This type II branchpoint is fundamentally different from the one in the initiator ubiquitin (see scheme in **Figure 5A**). In this case, Cdc48 can translocate both polypeptide branches through the pore, as demonstrated by the unfolding of fluorescent substrates located at either the N- or C-terminus of fusion proteins. Because Cdc48 translocates sequentially all ubiquitin molecules positioned between the initiator and substrate (proximal ubiquitins), it encounters many type II branchpoints during substrate processing, and at each of them likely unfolds both N- and C-terminal segments. The proximal ubiquitin molecules are all transiently unfolded but refold after translocation, while most substrates remain unfolded (**Figure 7B**, stage 7 to 8).

Because the D2 pore loops of Cdc48 surround a single polypeptide chain, it is possible that the ATPase translocates only one polypeptide branch at any given time and needs multiple rounds to unfold the entire substrate population. However, such a mechanism seems rather wasteful. An alternative is that Cdc48 moves several polypeptide chains simultaneously through its pore. Indeed, a structure of the related Vps4 ATPase with a circular peptide substrate indicates that a second strand can be accommodated (Han et al., 2019), and single-molecule experiments with the ClpB ATPase suggest that two strands can be processed at the same time (Avellaneda et al., 2020).

At the end of translocation, the entire polypeptide is released from the ATPase complex, because proximal ubiquitins and substrate translocate through the central pore, and distal ubiquitins move outside the pore (**Figure 7B**; stage 7). However, the unfolded, polyubiquitinated can re-bind to the ATPase complex, re-establish an initiation complex (similar to stage 2 in **Figure 7B**; not shown), and begin a new translocation cycle. The start of translocation seems to be rate-limiting as, according to our photocrosslinking experiments, the initiation state is most populated. For efficient transfer of the unfolded, polyubiquitinated substrate to the proteasome, the ubiquitin chain needs to be shortened by a deubiquitinase (DUB) to weaken its affinity for the ATPase complex. Indeed, Otu1, a DUB that binds to the N domains of Cdc48, can trim the ubiquitin chain (Bodnar and Rapoport, 2017b).

Our proteomics results indicate that the Cdc48 ATPase complex acts in vivo on the majority of polyubiquitinated proteins, showing little specificity for the actual substrate. This conclusion is consistent with our in vitro experiments and cryo-EM structure, which both show that the Cdc48 complex only recognizes ubiquitin molecules. Cofactors other than Ufd1 and Npl4, such as UBA-UBX proteins interacting with both Cdc48 and ubiquitin (Hänzelmann and Schindelin, 2017), could increase the binding constant for polyubiquitin chains, but since there are only a few of them, they cannot provide much substrate specificity. This raises the question as to how the Cdc48 complex, the proteasome, and the shuttling factors Rad23 and Dsk2 (Finley et al., 2012) divide their tasks, as they all seem to primarily recognize the ubiquitin chain, with little or no specificity for the actual substrate. In addition, the Cdc48/UN complex and the 26S proteasome require ubiquitin chains of about the same length (5 versus 4 ubiquitins (Bodnar and Rapoport, 2017b; Thrower et al., 2000)). So far, it has been assumed that Cdc48 is the most upstream component, then delivers substrates to the shuttling factors, which in turn bring them to the proteasome for degradation. However, with our new data, one would have to assume that Cdc48 unfolds every protein that carries a sufficiently long ubiquitin chain, a scenario that seems rather wasteful. An alternative is that the proteasome has the first pick (**Figure 7C**); only if a polyubiquitinated protein cannot be degraded would Cdc48 have a chance. In this model, the proteasome would act both upstream and downstream of the Cdc48 complex. Shuttling factors would transfer polyubiquitinated proteins back and forth between the Cdc48 complex and the proteasome, regardless of whether they are folded or unfolded (**Figure 7C**). They would pick up these proteins from the initiation state of the Cdc48 ATPase, the most populated state during substrate processing. Transfer between the Cdc48 complex and the 26S proteasome would be facilitated by DUB-mediated shortening of the ubiquitin chain. This model can now be tested with reconstituted systems containing purified Cdc48 complex, 26S proteasomes, shuttling factors, and possibly other Cdc48-interacting proteins.

## ACKNOWLEDGEMENTS

We thank the ICCB-Longwood Screening Facility for use of equipment. We thank Johannes Walter, Nicholas Bodnar, and Olga Kochenova for critical reading of the manuscript. This work was supported by a NIGMS grant (R01 GM052586) to T.A.R., by a research collaboration between J.R.E. and the Waters Corporation, by NIH grants (R01 CA233800 and R21 CA247671) to J.A.M., by financial support to J.A.M. from the Dana-Farber & Northeastern University Joint Program in Cancer Research, by an NIH/NIGMS grant (R01 GM132129) to J.A.P., and by an NIGMS grant (R01 GM067945) to S.P.G. Z.J. is a Howard Hughes Medical Institute Fellow of the Damon Runyon Cancer Research Foundation, DRG-2315-18. T.A.R. is a Howard Hughes Medical Institute Investigator.

## AUTHOR CONTRIBUTION

Z.J. and H.L. performed protein purifications, protein labeling, and all unfolding, binding, and photocrosslinking assays. D.P., T.E.W., and J.R.E. conducted the HDX MS analyses, J.A.P. and S.P.G. performed TMT labeling and MS analysis, and S.B.F. and J.A.M. analyzed crosslinked peptides by MS. T.A.R. supervised the project. Z.J. and T.A.R. wrote a draft of the manuscript.

## STAR * METHODS

### RESOURCE AVAILABILITY

#### Lead Contact

Further information and requests for resources and reagents should be directed to and will be fulfilled by the Lead Contact, Tom Rapoport.

#### Materials Availability

All unique/stable reagents generated in this study are available from the Lead Contact with a completed Materials Transfer Agreement.

#### Data and Code Availability

The HDX MS data have been deposited to the ProteomeXchange Consortium via the PRIDE partner repository with the dataset identifier PXD027639.

### EXPERIMENTAL MODEL AND SUBJECT DETAILS

#### Yeast strains and cultures

Plasmids encoding *Saccharomyces cerevisiae* Uba1 and Ubr1 were transformed into the INVSc1 yeast strain (Thermo). The yeast cells were grown in synthetic dropout (SD) medium for 24 h, and then switched to yeast culturing medium containing 2% galactose to induce protein expression. The cells were harvested after 24 h of the galactose induction.

The *pdr5*Δ strain was derived from BY4741 as described in (Yip et al., 2020). The pRS413(His) plasmid encoding either Npl4-FLAG or Ufd1-FLAG was transformed into the *pdr5*Δ strain. Positive clones on SD-Leu-His plates were pooled for subsequent experiments. Overnight cultures of yeast expressing either Npl4-FLAG or Ufd1-FLAG were inoculated into 80 ml of SD-Leu-His medium at an OD_600_ of 0.1. Cells were grown at 30°C until an OD_600_ of 0.7. The cells were then treated with 80 µM bortezomib (Selleckchem) or the same volume of DMSO and incubated at 30°C for another 4 h. 120 OD_600_ units of cells were spun down, and flash-frozen in liquid nitrogen.

#### Bacteria cultures

Bacterial expressing plasmids were transformed into *Escherichia coli* BL21 CodonPlus (DE3) RIPL cells (Agilent), unless stated otherwise. Bacterial strains were grown in Terrific Broth to an OD_600_ of 0.8. Protein expression was induced by addition of 0.1 mM isopropyl b-D-1-thiogalactopyranoside (IPTG), and then the incubation was continued at 16°C for 16 h.

p-Benzoyl-phenylalanine (Bpa) was incorporated into proteins by amber codon suppression in *Escherichia coli* BL21 (DE3) (New England BioLabs) harboring the plasmid pEVOL-pBpF(Chin et al., 2002). Cells were grown in Terrific Broth to an OD_600_ of 0.8. Protein expression was induced by the addition of 0.02% L-arabinose, 1 mM Bpa, and 0.2 mM IPTG, and the incubation was continued at 16°C for 16 h.

### METHOD DETAILS

#### Plasmids

For yeast experiments, the endogenous loci of the *Saccharomyces cerevisiae npl4* and *ufd1* genes – including their promoters, coding regions, and terminators – were amplified by polymerase chain reaction (PCR) and cloned into a yeast centromeric vector, pRS413(His3), using the NotI and XhoI restriction enzyme sites. A sequence encoding the FLAG tag (DYKDDDDK) was inserted at the C-terminus of each gene to express FLAG tagged Npl4 and Ufd1. *S. cerevisiae uba1* was cloned into the pRS426Gal1 vector with a His14-tag (HHHHSGHHHTGHHHHSGSHHH) and a TEV-protease cleavage site (ENLYFQG), as described in (Stein et al., 2014). *S. cerevisiae ubr1* gene was also cloned into pRS426Gal1(His14-TEV), using NotI and AscI sites, resulting the N-terminal sequence MSKHHHHSGHHHTGHHHHSGSHHHG-ENLYFQ-GAAA.

Wild-type Cdc48 and its variants were cloned into the pET28 vector using NotI and AscI sites, with a His6-tag and a TEV-protease cleavage site at the N-terminus. A sequence encoding the FLAG tag was added at the C-terminus of Cdc48, as appropriate. Cdc48^ND1^ contains residues 1-480 of wild-type Cdc48. Cdc48ΔD1Loops contains internal deletions of residues 286-290 and residues 325-330 of wild-type Cdc48. All Ufd1 variants were cloned into the pK27 vector with an N-terminal His14-SUMO (small ubiquitin-like modifier) tag. A sequence encoding the hemagglutinin (HA)-tag (YPYDVPDYA), or a streptavidin-binding protein (SBP) tag was inserted between SUMO and Ufd1, where indicated. Ufd1ΔUT3 contains a truncation of the first 200 residues of Ufd1. Ufd1^TEV^ has inserted a TEV-protease cleavage site (ENLYFQG) between the residues 209 and 210 of Ufd1. All Npl4 variants were cloned into the pET21 vector using NdeI and AscI sites, with a C-terminal His6-tag, FLAG tag, or FLAG-His6 tag. Internal deletions and insertions as well as point mutations were generated by overlapping PCR.

Wild-type human ubiquitin (hUb) and the I3C mutant were cloned into the pK27(His14-SUMO) vector using the Gibson assembly method. A sequence encoding a single Ala residue was inserted between the SUMO tag and the ubiquitin gene to allow for cleavage of the SUMO tag by Ulp1. All N-end rule degron fusions with a fluorescent substrate were constructed by overlapping PCR, and then cloned into the pK27(His14-SUMO) vector. The fluorescent substrates used in this study include the cysteine-free moxDendra2 (Dendra), the lysine-less super-folder GFP (sfGFP; a gift from Dirk Goerlich), the sfGFP-GGGSGGGSGGGS-Dendra fusion, and mEos3.2 (Eos; as described in (Bodnar and Rapoport, 2017b)). The sequence of the N-end rule degron is as follows (note that an N-terminal arginine is generated after SUMO cleavage): RHGSGCGAWLLPVSLVKRKTTLAPNTQTASPPSYRALADSLMQ. For substrates used in ubiquitin modification by a maleimide-conjugated fluorescent dye, the cysteine in the N-end rule degron was mutated to serine. All ubiquitin fusions with fluorescent proteins were constructed by overlapping PCR and cloned into the pET28(His6-FLAG) vector using the BamHI and AscI sites.

The bacterial expression plasmid encoding *S. cerevisiae* Ubc2 has been described in (Bodnar and Rapoport, 2017b). Plasmid encoding mouse Ube1 was a gift from Jorge Eduardo Azevedo (Addgene plasmid # 32534). The coding region of gp78^RING^-Ube2g2 was constructed as described previously (Blythe et al., 2017) and cloned into the pET28(His6-TEV) vector using NotI and AscI sites. The gp78^RING^-Ube2g2 fusion protein contains the RING domain of human gp78 (residue 322-393) and human Ube2g2 with the linker sequence GTGSH in between. The gene for human gp78 was a gift from Allan Weissman (Addgene plasmid # 37375). The gene for human Ube2g2 was a gift from Wade Harper (Addgene plasmid # 15791). The trypsin-resistant tandem ubiquitin binding entity (TR-TUBE) was cloned into pET28 vector using the NotI and AscI sites, resulting in the N-terminal sequence MGHHHHHHGSGENLYFQGAAACDI. The gene for TR-TUBE was a gift from Yasushi Saeki (Addgene plasmid # 110313). The pEVOL-pBpF plasmid used to produce Bpa-incorporated proteins was a gift from Peter Schultz (Addgene plasmid # 31190).

#### Immunoblotting and antibodies

Antibodies used in this study were: anti-Cdc48 (MyBioSource, MBS423348, 1:500), anti-ubiquitin (Santa Cruz Biotechnology, clone P4D1, 1:200), anti-FLAG (Sigma, clone M2, 1:1000), anti-HA (Roche, clone 12CA5, 1:1000), anti-ubiquitin K48-specific (Cell Signaling Technology, clone D9D5, 1:1000), anti-PGK1 (Abcam, clone 22C5D8, 1:1000), anti-SBP-tag (Millipore, clone 20, 1:1000), donkey anti-mouse IgG DyLight 800 conjugated (ThermoFisher, 1:5000), donkey anti-mouse IgG DyLight 680 conjugated (ThermoFisher, 1:5000), donkey anti-rabbit IgG DyLight 800 conjugated (ThermoFisher, 1:5000), donkey anti-goat IgG H&L horseradish peroxidase (HRP)-conjugated (Abcam, ab97110, 1:5000). The substrate for HRP conjugated secondary antibodies was Western Lighting Ultra (Perkin Elmer, NEL111001EA).

#### Protein purifications

All purified proteins were snap-frozen in size-exclusion chromatography (SEC) buffer (50 mM HEPES, pH 7.4, 150 mM NaCl, 5 mM MgCl_2_, and 0.5 mM tris(2-carboxyethyl)phosphine (TCEP)), except Cdc48^2Cys^, for which TCEP was omitted.

Cdc48, untagged Ufd1/Npl4 (UN), and the UN complexes harboring the groove mutants of Npl4 were expressed and purified as previously described (Twomey et al., 2019). Bacterial cells expressing Cdc48^2Cys^ were harvested by centrifugation at 5000 x g for 10 min and resuspended in wash buffer (50 mM Tris-HCl, pH 8, 320 mM NaCl, 5 mM MgCl_2_, 10 mM imidazole, 0.5 mM ATP) supplemented with phenylmethylsulfonyl fluoride (PMSF; 1 mM), a protease inhibitor cocktail, and DNase I (5 µg/ml). The cells were lysed by sonication. Lysates were cleared by ultracentrifugation in a Ti-45 rotor (Beckman) at 40,000 rpm for 30 min at 4°C. The supernatants were incubated with Ni-NTA resin that was pre-equilibrated with wash buffer, for 60 min at 4°C. The resin was washed three times with 30 column volumes of wash buffer. Proteins were eluted with elution buffer (50 mM Tris-HCl, pH 8, 150 mM NaCl, 5 mM MgCl_2_, 400 mM imidazole), and the eluates were diluted to about 800 nM and treated with 10 µM of the oxidant 4,4′-dipyridyl disulfide (Sigma) at 30°C for 30 min. After incubation, the reaction mixture was dialyzed against 50 mM HEPES, pH 7.5, 150 mM NaCl, 5 mM MgCl_2_ at 4°C overnight before snap-freezing.

Individual Ufd1 proteins were purified by Ni-NTA resin as described above. The SUMO protease Ulp1 was added to the eluted Ufd1 protein and dialyzed against wash buffer containing 10 mM imidazole. The Ulp1-treated samples were incubated with Ni-NTA resin to remove the His14-SUMO tag, and the unbound proteins were concentrated and loaded onto a Superdex 200 Increase column equilibrated with SEC buffer. Npl4 proteins were purified similarly by Ni-NTA and Superdex 200 Increase chromatography. Proteins with incorporated Bpa were purified in the same way as their parental counterparts.

Fluorescent substrates (Dendra, sfGFP, or Eos) containing the N-end rule degron and His14-SUMO tag were purified by Ni-NTA resin followed by the SUMO tag cleavage, SUMO tag removal with Ni-NTA resin, and gel filtration, similarly to the purification of the Ufd1 protein. Fluorescent substrates with ubiquitin fusions were purified similarly to the Npl4 protein by Ni-NTA resin and gel filtration.

His14-Uba1 was expressed in yeast cells and purified by Ni-NTA, ion-exchange, and size-exclusion chromatography, as described in (Stein et al., 2014). His14-Ubr1 was expressed and purified by Ni-NTA in the same way as His14-Uba1. After elution from the Ni-NTA resin, the protein was then buffer-exchanged into SEC buffer, concentrated, and snap-frozen. Ubc2 was expressed and purified as previously described (Twomey et al., 2019). *Mus musculus* Ube1, the gp78^RING^-Ube2g2 fusion, His14-SUMO-hUb, His14-SUMO-hUb^I3C^ and TR-TUBE were purified by Ni-NTA and size-exclusion chromatography. An additional step of SUMO cleavage and removal was performed prior to the gel filtration to purify wild-type hUb and the I3C mutant. Note that an extra alanine reside was left at the N-terminus of the purified hUb and hUb^I3C^ after SUMO cleavage. *S. cerevisiae* ubiquitin was purchased from Boston Biochem.

#### Dye labeling of substrates containing the N-end rule degron

The purified substrates were reduced with 10 mM TCEP and then incubated with a 3-fold molar excess of maleimide-conjugated DyLight dyes (Thermo). The reactions were kept in the dark at room temperature for 2 h before quenching with 20 mM dithiothreitol (DTT). The unreacted free dyes were removed by Dye Removal columns (Thermo, #22858).

#### Photoconversion of substrates containing Dendra or Eos

The purified substrate proteins (4∼8 mg/ml) were placed in a 200-µl PCR tube in an ice bath. A long-wavelength UV flashlight (395-410 nm, DULEX) was positioned 5 cm above the tube, and the sample was irradiated for 1 h, with occasional mixing.

#### Ubiquitination of substrates

Ubiquitination of the substrates containing an N-end rule degron was carried out as previously described (Twomey et al., 2019), with some modifications. Substrate (5 µM) was incubated with *S. cerevisiae* ubiquitin or purified human ubiquitin (250 µM), Uba1 (800 nM), Ubc2 (4.63 µM), Ubr1 (800 nM), and ATP (10 mM) for 60 min at 30°C in ubiquitination buffer (50 mM Tris pH 8.0, 150 mM NaCl, 10 mM MgCl_2_, 1 mM DTT). The samples were concentrated and loaded onto a Superdex 200 column in SEC buffer. After analysis by SDS-PAGE, fractions containing the desired ubiquitin chain lengths were pooled and snap-frozen. The concentration of the pooled polyubiquitinated substrate was determined with a Synergy Neo2 Multi-mode reader (BioTek), using the non-ubiquitinated substrates as standards. The majority of the final product contained polyubiquitin chains of 10-25 ubiquitin molecules. The [Dendra-Ub](n)-Dendra substrate in Figure 3F was generated in a reaction containing 4 µM dye-labeled, photo-converted Dendra fusion with N-end rule degron and 50 µM Dendra-hUb fusion.

Ubiquitination of the ubiquitin fusion substrates were performed as described in (Blythe et al., 2017) with some modifications. 10 µM the ubiquitin-fusion substrate was incubated with 1 µM mouse Ube1, 20 µM gp78^RING^-Ube2g2, and 500 µM purified human ubiquitin in 20 mM HEPES, pH 7.4, 100 mM NaCl, 2 mM DTT, 10 mM ATP, and 10 mM MgCl_2_, and incubated at 37°C. The 500 µM ubiquitin and 10 mM ATP were added in small amounts every 30 min over the first 5 h. The reaction was then kept at 37°C overnight. The samples were incubated with Ultra HBC streptavidin agarose beads (Goldbio) or FLAG antibody M2 agarose resin (Sigma) for 60 min at 4°C. The resin was washed three times with 5 volumes of ubiquitination buffer and bound material eluted with 5 volume of ubiquitination buffer containing 2 mM biotin or 1 mg/ml 3xFLAG peptide. Subsequent gel filtration, SDS-PAGE analysis, and concentration determination were performed as described above. To generate polyubiquitin chains containing Dendra-hUb^K48R^, the ubiquitination reaction was first carried out with 1 µM mouse Ube1, 20 µM gp78^RING^-Ube2g2, and 500 µM wild-type human ubiquitin at 30°C for 5 h. 20 µM Dendra-hUb^K48R^ was then added to the reaction prior to the overnight incubation.

#### HDX MS measurements

Photoconverted, polyubiquitinated SBP-Dendra substrates were bound to streptavidin agarose resin (Thermo) and incubated at 4°C with Ufd1, Npl4, and Cdc48 at a 1:1:1 molar ratio in assembly buffer (50 mM HEPES, pH 7.5, 100 mM NaCl, 5 mM MgCl_2_, 1 mM DTT). Note that no nucleotides were present during the incubation. The resin was washed to remove unbound Cdc48/UN complex. The beads were washed once with four bead volumes of assembly buffer and then washed twice with four volumes of HDX equilibration buffer (20 mM Tris, 150 mM NaCl, 5 mM MgCl_2_, 0.5 mM TCEP, 100% H_2_O, pH 7.5). Bound protein was eluted with the HDX equilibration buffer containing 2 mM biotin, and concentrated. The final sample contained about 15 µM Cdc48, 16 µM UN, and 36 µM Ub(n)-Dendra. As controls, the Cdc48/UN complex and photoconverted, polyubiquitinated SBP-Dendra were also buffer-exchanged separately into HDX equilibration buffer. 8 µM of Cdc48/UN, Ub(n)-Dendra alone, or the Cdc48/UN in complex with Ub(n)-Dendra were mixed with 10 mM of ADP or ATP and incubated at 30°C for 30 min, and then kept on ice until HDX began.

Deuterium labeling and measurement were performed as previously described (Twomey et al., 2019). To initiate HDX, 1.0 µl of each protein sample was diluted at 20°C with 18 μl labeling buffer (20 mM Tris-HCl, 150 mM NaCl, 5 mM MgCl_2_, and 0.5 mM TCEP, 99.9% D2O, pD 7.4). At each labeling time (5 s, 10 s, 1 min, 10 min, and 4 h), 19 μl of quench buffer was added (150 mM potassium phosphate pH 2.49, H_2_O). All subsequent steps were performed at 0 °C. Quenched samples were digested online with immobilized pepsin using a Waters UPLC instrument with HDX technology (Wales et al., 2008), desalted, and then eluted into a Waters Synapt XS mass spectrometer with a 5-35% gradient of water:acetonitrile over 10 min. Peptic peptides were identified using ProteinLynx Global Server (PLGS) 3.0.1 (Waters) and deuterium incorporation measured using DynamX 3.0 (Waters). The deuterium levels were not corrected for back exchange and are reported as relative (Wales and Engen, 2006). The error of measuring deuterium in this LC/MS setup was +/- 0.20 relative Da and differences were considered meaningful if they were larger than 0.50 Da. The recommended summary (Masson et al., 2019) of HDX MS experimental parameters, proteolytic maps for all proteins, and the numeric values used to create HDX-MS figures are provided in **Table S1**. The raw HDX MS data and expanded technical details of the HDX MS acquisition and data processing steps have been deposited to the ProteomeXchange Consortium via the PRIDE partner repository (Perez-Riverol et al., 2019) with the dataset identifier PXD027639.

#### Ubiquitin modification by a maleimide-conjugated fluorescent dye

A cysteine-free substrate containing the N-end rule degron and photoconverted Dendra was polyubiquitinated with purified hUb^I3C^, as described above, resulting in Ub^I3C^(n)-Dendra. 400 nM of Ub^I3C^(n)-Dendra were incubated with 400 nM Ufd1, 400 nM Npl4, and 1 µM full-length Cdc48 or Cdc48^ND1^ in SEC buffer containing 10 mM ADP for 40 min at 4°C. The samples were then supplemented with 100 µM of maleimide-conjugated DyLight 680 dye (Thermo), incubated for another 10 min, and quenched by the addition of 20 mM DTT. The samples were mixed with SDS sample buffer and analyzed by SDS-PAGE, followed by fluorescence scanning on an Odyssey imager (LI-COR) and Coomassie-blue staining.

#### Substrate unfolding assays

With the exceptions mentioned below, substrate unfolding experiments were performed as previously described (Twomey et al., 2019). Briefly, 400 nM of the polyubiquitinated, photoconverted Dendra or Eos proteins were mixed with 400 nM UN variants and 400 nM Cdc48 variants in 50 mM HEPES pH 7.5, 150 mM NaCl, 10 mM MgCl_2_, 0.5 mM TCEP, and 0.5 mg/ml protease-free bovine serum albumin (BSA). After a 10-min pre-incubation at 30°C, an ATP regeneration mixture was added (10 mM ATP, 20 mM phosphocreatine, 100 µg/ml creatine kinase), and the fluorescence (excitation, 540 nm; emission, 580 nm; gain, 80 to 100) was measured at 15-s intervals for 30 min, using a Synergy Neo2 Multi-mode reader (BioTek). Fluorescence of the unfolding reactions in Figures 4B, S2F, and S5A were measured at 30-s intervals for 30 min, using FlexStation 3 Microplate Reader (Molecular Devices).

The unfolding reactions in Figure 3E contained no TCEP. 800 nM of the oxidized Cdc48^2Cys^ was pre-treated with 20 mM DTT to generate the reduced form of Cdc48^2Cys^. For protease treatment of polyubiquitinated substrates, 2 µM of substrates were incubated with 25 µM TEV protease (Figure 3F) or 9 µM Ulp1 (Figure S2F) in SEC buffer containing 1 mM DTT at 30°C for 1 h, before being added to the unfolding reactions. Unfolding reactions in Figure 5C contained 800 nM Ufd1/Npl4 and 800 nM Cdc48, the ones in Figure S5F contained 800 nM substrate, 800 nM Ufd1/Npl4, and the indicated concentrations of Cdc48.

Unfolding assays performed in conjunction with pull-downs (Figures 6F, S5A-E) were described in the section “in vitro pull-down experiments”. Unfolding reactions in Figure S5C-D were scanned in dual fluorescence mode with an additional green fluorescence channel (excitation 2,488 nm; emission 2, 525 nm; gain 2, 80). Unfolding assays in conjunction with photocrosslinking (Figures 4B and S7A) were described in the section “photocrosslinking experiments”. Unfolding reactions in Figure S7E were carried out with 400 nM Ub^I3C^(n)-Dendra, 400 nM HA-Ufd1^TEV^, 400 nM Npl4-FLAG, 1 µM Cdc48, and 10 mM of the indicated nucleotides. Protease-free BSA was omitted to avoid competition of BSA with the maleimide-conjugated DyLight 680 dye. After unfolding, the samples were incubated on ice for 1 h in the presence or absence of 1 µM TEV protease, prior to the addition of the maleimide-conjugated dye. Subsequent ubiquitin modification assays were performed as described above.

For data acquired on Synergy Neo2 Multi-mode reader (BioTek), the relative fluorescence at time t was calculated as (fluorescence at t) / (fluorescence at t0). For each experiment performed on FlexStation 3 Microplate Reader (Molecular Devices), an empty well was included to determine background fluorescence. The relative fluorescence at time t was calculated as [(fluorescence at t) – (background fluorescence at t)] / [(fluorescence at t0) – (background fluorescence at t0)]. The calculated relative fluorescence was plotted against time using Prism software (GraphPad). A linear fit was performed with data points within the first 2 min to calculate the initial velocities. These rates were normalized to that of the wild-type sample in the same experiment. Inhibition of substrate unfolding was determined by the reduction of the initial unfolding velocity compared to the wild-type sample. See **Table S2** for the raw data of all substrate unfolding assays.

#### In vitro pull-down experiments

For all in-vitro pull-down experiments, with the exceptions mentioned below, 1 µM polyubiquitinated substrate was mixed with 1 µM Cdc48, 1 µM Ufd1, and/or 1 µM Npl4 in binding buffer (50 mM Tris-HCl, pH 8, 150 mM NaCl, 10 mM MgCl_2_, and 1 mM DTT) supplemented with 10 mM of the desired nucleotide. 50 µl of such a protein mixture were then incubated with 8 µl pre-equilibrated FLAG antibody M2 agarose beads (Sigma) or streptavidin agarose beads (Thermo) at 4°C for 1 h. The beads were then washed three times with binding buffer containing the desired nucleotide. Bead-bound proteins were eluted with 25 µl of binding buffer supplemented with 0.05% Tween-20 and either 0.2 mg/ml 3xFLAG peptide (Bimake) or 2 mM biotin (Sigma). The eluted samples were subjected to SDS-PAGE, followed by fluorescence scanning on an Odyssey imager (LI-COR) and Coomassie-blue staining. When oxidized Cdc48^2Cys^ was used for pull-downs, all proteins were buffer-exchanged into non-reducing SEC buffer, and the binding reactions were carried out in a non-reducing binding buffer (50 mM Tris-HCl, pH 8, 150 mM NaCl, and 10 mM MgCl_2_).

For the substrate-dissociation assay shown in Figures 6C-D, substrate unfolding reactions were performed with 200 nM substrate, 1 µM Cdc48, 1 µM Ufd1^TEV^, 1 µM Npl4-FLAG, and 10 mM of ADP or ATP. 50 µl of each reaction were then incubated with 5 µl pre-equilibrated FLAG antibody M2 agarose beads (Sigma) at 4°C for 1 h. After washing with nucleotide-free binding buffer, the beads were resuspended in 50 µl unfolding buffer containing 4.5 µM TEV protease and 10 mM of the desired nucleotide, and the incubation was continued at room temperature for another 2 h. After three washes with the binding buffer supplemented with appropriate nucleotides, the bound proteins were eluted and analyzed as described above.

For the substrate exchange experiment shown in Figure 6E, the unfolding reactions were performed with 1 µM DyLight 800-labeled substrate, 2 µM Cdc48-FLAG, 4 µM SBP-Ufd1, 4 µM Npl4, and 10 mM of ADP or ATP. 50 µl of each reaction were then incubated with 5 µl pre-equilibrated streptavidin agarose beads (Thermo) at 4°C for 1 h. After three washes with assembly buffer (50 mM HEPES, pH 7.5, 100 mM NaCl, 5 mM MgCl_2_, 1 mM DTT), bound proteins were eluted by 2 mM biotin in assembly buffer. The eluted proteins were mixed with 1 µM DyLight 680-labeled substrate and 10 mM of the desired nucleotide for a second unfolding reaction. The samples were then incubated with 5 µl pre-equilibrated FLAG antibody M2 agarose beads (Sigma) for 1 h at 4°C, followed by washing, elution, and SDS-PAGE, as described above.

#### Photocrosslinking experiments

The photocrosslinking experiments in Figures S2D, S3D, and S3I were performed as described (Twomey et al., 2019), with some modifications. Briefly, the reaction components included Cdc48_ND1^D324Bpa^-FLAG, Cdc48^D324Bpa^-FLAG, or Cdc48^D602Bpa^-FLAG (200 nM), wild-type Ufd1 or Ufd1ΔUT3 (500 nM), wild-type or mutant Npl4 (500 nM), and dye-labeled polyubiquitinated sfGFP (1 µM) in reaction buffer (50 mM HEPES pH 7.5, 150 mM NaCl, 10 mM MgCl_2_, 0.5 mM TCEP, 0.5 mg/ml protease-free BSA) supplemented with 10 mM of ADP. The reactions were assembled on ice, incubated at 30°C for 10 min, and transferred to individual PCR tubes. A long-wave UV lamp (Blak-Ray) was positioned 5 cm above the tubes, and the samples were irradiated for 30 min. To prevent overheating, an ice-cold metal block was placed in contact with the bottom of the PCR tubes. After irradiation, the samples were diluted 10-fold in dissociation buffer (50 mM Tris-HCl, pH 8, 800 mM NaCl, 1% (v/v) Triton X-100, 1 mM EDTA, 0.5 mM DTT) and incubated at room temperature for 5 min. The samples were then applied to 8 µl FLAG antibody M2 magnetic beads (Sigma) equilibrated in dissociation buffer for 1 h at room temperature. The beads were washed three times, and bound protein was eluted in 50 mM HEPES pH 7.5, 150 mM NaCl, 0.5 mM TCEP, and 0.2 mg/ml 3xFlag peptide (Bimake). The eluted material was subjected to SDS-PAGE and the gel scanned on an Odyssey imager (LI-COR). The gel was then transferred to a nitrocellulose membrane and analyzed by immunoblotting with Cdc48 antibodies on an Amersham Imager 600 RGB (Cytiva).

The photocrosslinking experiment in Figure 4B was performed with 1.25 µM Cdc48, 1.25 µM SBP-Ufd1, 1.25 µM FLAG-tagged Npl4 Bpa mutant, and 1.25 µM dye-labeled, photoconverted, and polyubiquitinated Dendra, in 50 mM HEPES, pH 7.5, 100 mM NaCl, 5 mM MgCl_2_, and 1 mM DTT. 16 µl of the protein mixture were then diluted with 29 µl of unfolding buffer (50 mM HEPES, pH 7.5, 150 mM NaCl, 10 mM MgCl_2_, 0.5 mg/ml protease-free BSA). After a 10-min pre-incubation at 30°C, 5 µl of ATP (100 mM) was added and the fluorescence (excitation, 540 nm; emission, 580 nm) was measured at 30-s intervals for 30 min, using a FlexStation 3 Microplate Reader (Molecular Devices). Both crosslinked and non-crosslinked unfolding reactions were analyzed by SDS-PAGE. The non-crosslinked sample was also subjected to UV irradiation after substrate unfolding, and analyzed by SDS-PAGE side-by-side, as shown in Figure S4B.

For photocrosslinking experiment followed by MS (Figures S7A-D), an unfolding assay was first performed with 400 nM Cdc48^D602Bpa^-FLAG, 600 nM Ufd1/Npl4, 400 nM dye-labeled, photo-converted, and polyubiquitinated Dendra, and 10 mM ADP or ATP. Unfolding of Dendra was monitored at 15-s intervals for 30 min using a Synergy Neo2 Multi-mode reader (BioTek). The samples were then transferred to a PCR tube for UV irradiation. The Cdc48^D602Bpa^-FLAG protein was isolated by FLAG antibody M2 magnetic beads (Sigma) and bound material eluted with 0.2 mg/ml single FLAG peptide (Sigma-Aldrich) in 50 mM Tris-HCl, pH 8, 150 mM NaCl, 0.05% NP-40, 1 mM EDTA, 10% glycerol, and protease inhibitor cocktail. The eluted material was subjected to both SDS-PAGE and MS analysis.

#### MS of photocrosslinked proteins

The analysis of crosslinked peptides was performed by nano-liquid chromatography and tandem MS, as previously described (Twomey et al., 2019), with minor modifications. FLAG peptide eluates were diluted 1:1 with 100 mM ammonium bicarbonate, denatured with 0.1% Rapigest (Waters Corporation, Milford, MA), reduced with 10 mM DTT for 30 min at 56 °C, cooled for 5 min at room temperature, alkylated with 22.5 mM iodoacetamide for 30 min at room temperature protected from light, and then digested with trypsin overnight at 37°C. Rapigest was cleaved by adding trifluoroacetic acid to a final concentration of 1% and incubating for an additional 30 min at 37°C. After centrifugation to remove Rapigest by-products, peptides in the supernatant were desalted using C18 (SOLA-RP, ThermoFisher Scientific, Madison, WI). C18 eluates were dried by vacuum centrifugation, and residual detergent was removed using magnetic beads (Hughes et al., 2014).

Peptides were analyzed by nanoLC-MS as described (Ficarro et al., 2009) using a NanoAcquity UPLC system (Waters Corporation) interfaced to a QExactive HF mass spectrometer (ThermoFisher Scientific). Peptides were injected onto a self-packed pre-column (100 µm I.D. packed with 4 cm POROS 10R2, Applied Biosystems, Framingham, MA), resolved on an analytical column (30 µm I.D. packed with 50 cm 5 µm Monitor C18, Orochem, Naperville, IL), and introduced to the mass spectrometer via ESI (spray voltage = 4 kV). The mass spectrometer was programmed to operate in data dependent mode, such that the 15 most abundant precursor ions in each MS scan (*m/z* 300-2000, 120K resolution, target=1E6) were subjected to MS/MS (target value=5E4, max fill time=50 ms, isolation width=1.6 Da, resolution=15K, collision energy=30%).

Cross-linked peptides were identified using CrossFinder version 1.4 (Mueller-Planitz, 2015) to search against a customized protein sequence database consisting of Cdc48, Ufd1, Npl4, ubiquitin, and the fusion protein of N-end-rule degron and the fluorescent protein Dendra.

#### Immunoprecipitation of yeast lysates

Immunoprecipitation (IP) from lysates of *S. cerevisiae* cells was performed as described in (Tsuchiya et al., 2017) with some modifications. Briefly, a cell pellet corresponding to 120 OD_600_ was resuspended in 800 µl of lysis buffer (50 mM Tris-HCl, pH 7.5, 100 mM NaCl, 10% glycerol, 10 µM bortezomib, 10 mM iodoacetomide, 1x Protease Inhibitor Cocktail (Roche)) and mixed with 1 ml of acid-washed glass beads. The cells were lysed using a Mini-BeadBeater 96 (BIOSPEC) at 2,400 rpm for 4.5 min. The cell lysate was then spun at 500 g for 3 min at 4°C to remove unbroken cells and debris. The supernatant was supplemented with 1% TX-100 and incubated on ice for 30 min, before spinning at 20,000 g for 20 min. A small aliquot of the cleared lysate was used for immunoblots with K48-specific ubiquitin antibodies and anti-PGK1 antibodies. The protein concentration in the cleared lysate was determined with a Protein Assay Dye Reagent (BioRad). One half of the lysate containing ∼2.5 mg of proteins was incubated with 25 µl FLAG antibody agarose resin (Sigma, M2) that was pre-equilibrated with lysis buffer containing 1% TX-100. The other half of the lysate was incubated with 5 µg of biotinylated, K48-specific TUBE (LifeSensors, UM307) and 25 µl pre-equilibrated streptavidin agarose beads (ThermoFisher). After one hour incubation at 4°C, the beads were washed three times with 1 ml of the lysis buffer supplemented with 1% TX-100. Bound material was eluted with 4 bead-volumes of elution buffer (50 mM HEPES, pH 7.4, 150 mM NaCl, 5 mM MgCl_2_, 0.5 mM TCEP, 0.1% TX-100) supplemented with 1 mg/ml TR-TUBE. 20 µl of the total eluate was examined by SDS-PAGE, silver stain, and immunoblotting using K48-specific ubiquitin antibodies. The other 80 µl of the eluate were precipitated by adding 20 µl of 100% trichloroacetic acid (TCA) (Sigma). After 30-min incubation on ice, the samples were centrifuged at 14,000 rpm for 15 min at 4°C. The pellet was sequentially washed with 1 ml of ice-cold acetone and 1 ml of cold methanol. The protein pellet was then resuspended and digested with trypsin for MS analysis.

#### TMT labeling

TMTpro reagents (0.8 mg) were dissolved in anhydrous acetonitrile (40 μl) of which 7 μl were added to the peptides (50 µg). Acetonitrile (13 μl) was added to achieve a final concentration of approximately 30% (v/v). Following incubation at room temperature for 1 h, the reaction was quenched with hydroxylamine at a final concentration of 0.3% (v/v). Equal amounts of all TMTpro-labeled samples were pooled, dried under vacuum in a SpeedVac, and subjected to a C18 solid-phase extraction (SPE) column with a capacity of 100 mg (Sep-Pak, Waters).

#### Liquid chromatography and tandem MS

MS data were collected on an Orbitrap Eclipse mass spectrometer coupled to a Proxeon NanoLC-1200 UHPLC. The 100 µm capillary column was packed with 35 cm of Accucore 150 resin (2.6 μm, 150Å; ThermoFisher Scientific). Data were acquired for 180 min for a total of 4 injections of the unfractionated sample. The scan sequence began with an MS1 spectrum (Orbitrap analysis, resolution 120,000, 400−1500 Th, automatic gain control (AGC) target was set to “standard”, maximum injection time was set to “auto”). The number of data dependent scans was set to 20. MS2 analysis consisted of higher-energy collision-activated dissociation (HCD), the Orbitrap resolution was set at 50,000, the isolation window was 0.7 Da, automatic gain control (AGC) was set at 300%, and HCD collision energy was 37%. All data were acquired with FAIMS with the dispersion voltage (DV) set at 5000V. Injections differed in the compensation voltages (CVs) used. One sample was subjected to CV= -40, -60, and -80V, a second sample with -50 and -70V and two samples with -45, -55, -65, and -75V. In one of the samples with the four CVs, the precursor priority was set from least intense to most intense.

Spectra were converted to mzXML via MSconvert (Chambers et al., 2012). Database searching included all entries from the yeast UniProt Database (downloaded: August 2020). The database was concatenated with one composed of all protein sequences for that database in the reversed order. Searches were performed using a 50-ppm precursor ion tolerance for total protein level profiling. The product ion tolerance was set to 0.02 Da. These wide mass tolerance windows were chosen to maximize sensitivity in conjunction with Comet searches and linear discriminant analysis (Beausoleil et al., 2006; Huttlin et al., 2010). TMTpro labels on lysine residues and peptide N-termini (+304.207 Da), as well as carbamidomethylation of cysteine residues (+57.021 Da) were set as static modifications, while oxidation of methionine residues (+15.995 Da) was set as a variable modification. Peptide-spectrum matches (PSMs) were adjusted to a 1% false discovery rate (FDR) (Elias and Gygi, 2007; 2010). PSM filtering was performed using a linear discriminant analysis, as described previously (Huttlin et al., 2010) and then assembled further to a final protein-level FDR of 1% (Elias and Gygi, 2007). Proteins were quantified by summing reporter ion counts across all matching PSMs, also as described previously (McAlister et al., 2012). Reporter ion intensities were adjusted to correct for the isotopic impurities of the different TMTpro reagents according to manufacturer specifications. The signal-to-noise (S/N) measurements of peptides assigned to each protein were summed and these values were normalized so that the sum of the signal for all proteins in each channel was equivalent to account for equal protein loading. Finally, each protein abundance measurement was scaled, such that the summed signal-to-noise for that protein across all channels equals 100, thereby generating a relative abundance measurement.

For each substrate protein detected, the ratio of its abundance in the FLAG IP versus K48 IP from the same cell lysate was calculated. The two ratios calculated from cell lysates under the same condition were averaged as the Cdc48/K48 ratio. The averaged Cdc48/K48 ratios of all substrate proteins from four different conditions – Npl4-FLAG/DMSO, Ufd1-FLAG/DMSO, Npl4-FLAG/bortezomib, and Ufd1-FLAG/bortezomib – were then plotted as pairwise comparisons in a logarithmic scale. After confirming the overall distributions of Cdc48/K48 ratio across all the four conditions were very similar, the Cdc48/K48 ratios of each substrate protein were further averaged. The resulting Cdc48/K48 ratios were plotted in a logarithmic scale to generate a distribution plot. All graphs were generated using Prism software (GraphPad).

### QUANTIFICATION AND STATISTICAL ANALYSIS

Quantifications of fluorescence scanning gels were carried out using the ImageStudio software (LI-COR). For each lane, a rectangle box was selected to determine total intensity of a band or smear of signal. The rectangle boxes for all lanes on the same gel were kept with similar box sizes. For each gel, an additional rectangle box with similar box size were drawn over an empty or non-signal region to determine background intensity. Signal intensity of each lane was calculated as (total intensity – box size * background intensity / background box size). The resulting signal intensity was normalized to a designated lane to calculate the relative signal intensity.

Band intensities on Coomassie blue-stained gels were quantified using ImageJ (NIH). Background subtraction and normalization were performed the same as fluorescence scanning gels described above.

### SUPPLEMENTAL INFORMATION

Supplemental information includes: Document S1. Figures S1-S7

Table S1. Hydrogen-deuterium exchange mass spectrometry data (related to Figures 2 and S6)

Table S2. Raw data of substrate unfolding assays (related to Figures 3-6, S2, S3, S5, and S7)

