## Supplemental Document S1 for "Translocation of polyubiquitinated protein substrates by the hexameric Cdc48 ATPase"

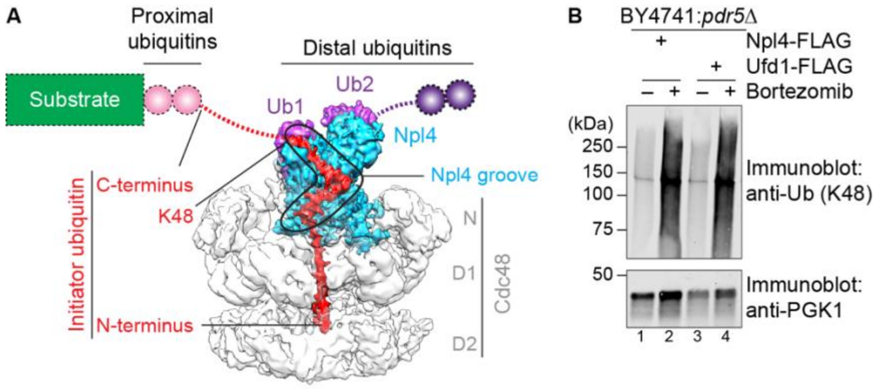


**Figure S1 (related to Figure 1). Cdc48 complex with bound ubiquitin molecules.**

**(A)** Cryo-EM map of the Cdc48 initiation complex (EMD-0665). Cdc48 is shown in grey, with the N, D1, and D2 domains indicated. The Npl4 cofactor is colored in blue, the unfolded initiator ubiquitin in red, and the folded ubiquitin molecules Ub1 and Ub2 in purple. The groove of Npl4 is highlighted. The invisible C-terminus of the initiator ubiquitin is indicated by a red dotted line, and invisible proximal and distal ubiquitins by pink and purple circles, respectively. Substrate is shown schematically as a green box.

**(B)** *S. cerevisiae* cells (strain BY4741) expressing Npl4-FLAG or Ufd1-FLAG were incubated in the absence or presence of the proteasome inhibitor bortezomib. Cell lysates were subjected to SDS-PAGE, followed by immunoblotting with K48-specific polyubiquitin antibodies. Blotting for phosphoglycerate kinase 1 (PGK1) served as a loading control.

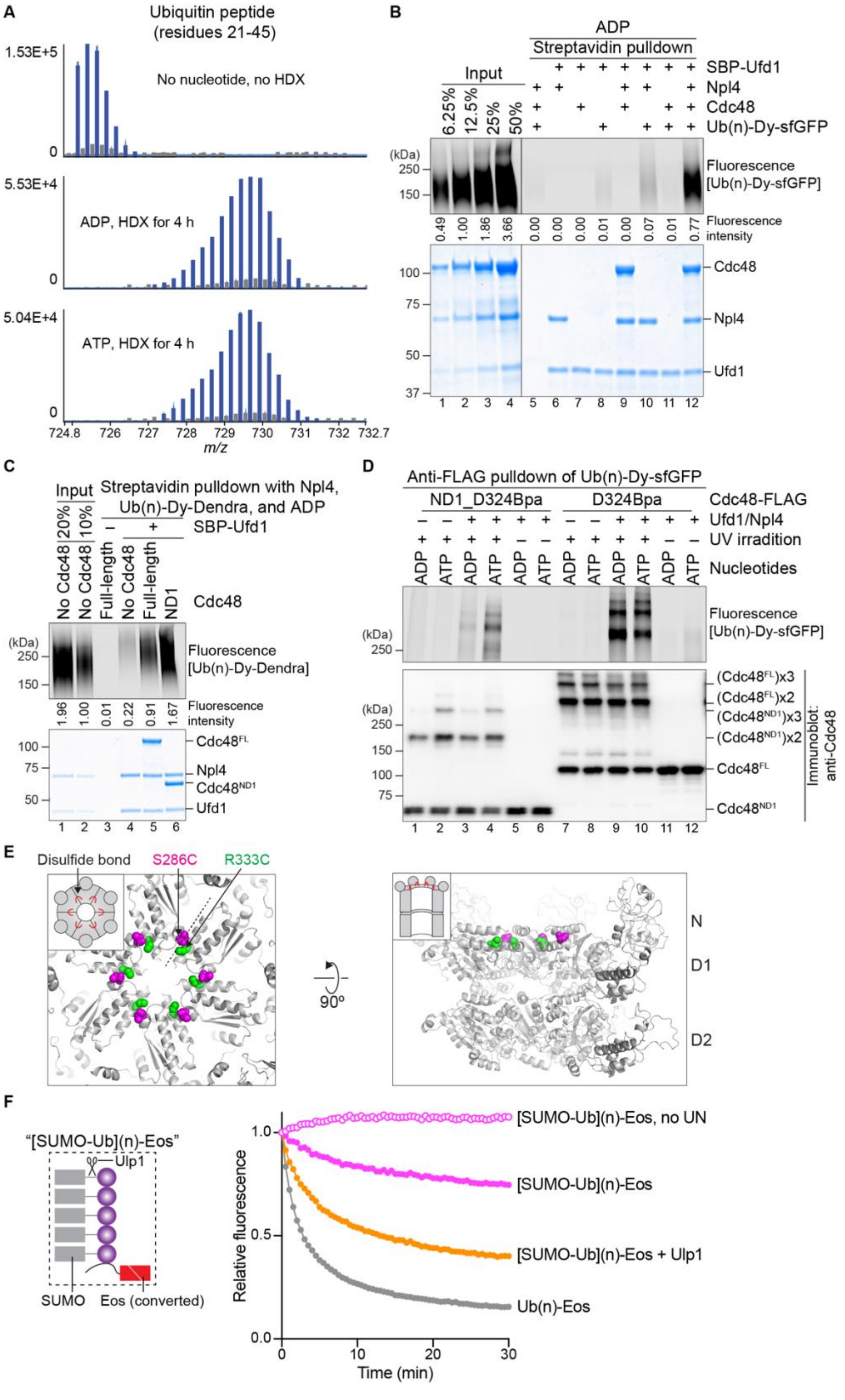

A
Ubiquitin peptide (residues 21-45)
1.53E+5
No nucleotide, no HDX
5.53E+4
ADP, HDX for 4 h
5.04E+4
ATP, HDX for 4 h
0
724.8 726 727 728 729 730 731 732 732.7
m/z
B
ADP
Streptavidin pulldown
Input
6.25% 12.5% 25% 50%
SBP-Ufd1
Npl4
Cdc48
Ub(n)-Dy-sfGFP
(kDa)
250
150
Fluorescence [Ub(n)-Dy-sfGFP]
0.49 1.00 1.86 3.66 0.00 0.00 0.00 0.01 0.00 0.07 0.01 0.77
Fluorescence intensity
100 75 50 37
Cdc48
Npl4
Ufd1
1 2 3 4 5 6 7 8 9 10 11 12
C
Input
20% 10%
Streptavidin pulldown with Npl4, Ub(n)-Dy-Dendra, and ADP
SBP-Ufd1
No Cdc48 No Cdc48 Full-length No Cdc48 Full-length ND1
Cdc48
(kDa)
250
150
Fluorescence [Ub(n)-Dy-Dendra]
1.96 1.00 0.01 0.22 0.91 1.67
Fluorescence intensity
100 75 50
Cdc48FL
Npl4
Cdc48ND1
Ufd1
1 2 3 4 5 6
D
Anti-FLAG pulldown of Ub(n)-Dy-sfGFP
ND1_D324Bpa
D324Bpa
Cdc48-FLAG
Ufd1/Npl4
UV irradition
Nucleotides
ADP ATP ADP ATP ADP ATP ADP ATP ADP ATP ADP ATP
Fluorescence [Ub(n)-Dy-sfGFP]
(kDa)
250
(Cdc48FL)x3
(Cdc48FL)x2
(Cdc48ND1)x3
(Cdc48ND1)x2
Cdc48FL
Cdc48ND1
Immunoblot: anti-Cdc48
250 150 100 75
1 2 3 4 5 6 7 8 9 10 11 12
E
Disulfide bond
S286C
R333C
90º
N
D1
D2
F
"[SUMO-Ub](n)-Eos"
Ulp1
SUMO
Eos (converted)
[SUMO-Ub](n)-Eos, no UN
[SUMO-Ub](n)-Eos
[SUMO-Ub](n)-Eos + Ulp1
Ub(n)-Eos
Relative fluorescence
1.0 0.5 0.0
0 10 20 30
Time (min)

**Figure S2. (related to Figures 2 and 3). Cdc48 enhances substrate recruitment by polypeptide insertion into the D1 ring.**

**(A)** Photoconverted Ub(n)-Dendra was incubated with Cdc48/UN complex in the presence of ADP or ATP and subsequently subjected to HDX for 4 hrs. A control was performed without deuteration. All samples were subjected to proteolytic cleavage, and the deuteration of ubiquitin peptides determined by MS. Blue bars show peptides that were assigned to the ubiquitin peptide comprising residues 21-45, and grey bars show unassigned peptides and peptides that were assigned to other protein segments.

**(B)** Ub(n)-sfGFP labeled with a fluorescent dye was incubated in the presence of ADP with different combinations of SBP-Ufd1, Npl4, and Cdc48. Streptavidin beads were added and bound material analyzed by SDS-PAGE, followed by fluorescence scanning (upper panel) and Coomassie-blue staining (lower panel). To evaluate the pull-down efficiency, different amounts of the input material were loaded (left four lanes).

**(C)** Dye-labeled Ub(n)-Dendra was incubated in the presence of ADP with Npl4, SBP-Ufd1, and full-length Cdc48 or Cdc48 containing only the N and D1 domains (ND1). Material bound to streptavidin beads was analyzed as in (B).

**(D)** Dye-labeled Ub(n)-sfGFP was incubated with Ufd1, Npl4, and either full-length Cdc48 ($Cdc48^{FL}$) or the ND1 mutant ($Cdc48^{ND1}$), both containing a photoreactive Bpa probe in the D1 ring (D324Bpa). The samples were irradiated, as indicated, and the Cdc48 complex was retrieved with FLAG antibody beads. Bound material was analyzed by fluorescence scanning (upper panel) or immunoblotting with Cdc48 antibodies (lower panel). Note that substrate not only crosslinks to monomeric $Cdc48^{FL}$ and $Cdc48^{ND1}$, but also to crosslinked dimers and trimers.

**(E)** A Cdc48 hexamer was generated in which all subunits are disulfide-crosslinked to their neighbors. Shown are top and side views of the D1 ring (left and right panel, respectively), with the Cys residues forming disulfides shown as red and green spheres. The dashed line indicates the interface between adjacent protomers. In the side view, the three Cdc48 subunits in the front were removed.

**(F)** A fusion of the N-end-rule degron with the fluorescent protein Eos was irradiated with UV light, causing photoconversion of Eos. A chain of SUMO-ubiquitin fusions was assembled on a lysine residue within the N-terminal degron, resulting in [SUMO-Ub](n)-Eos (see scheme on the left). After incubation with or without the SUMO protease Ulp1, unfolding of Eos was tested with Cdc48 in the absence or presence of UN.

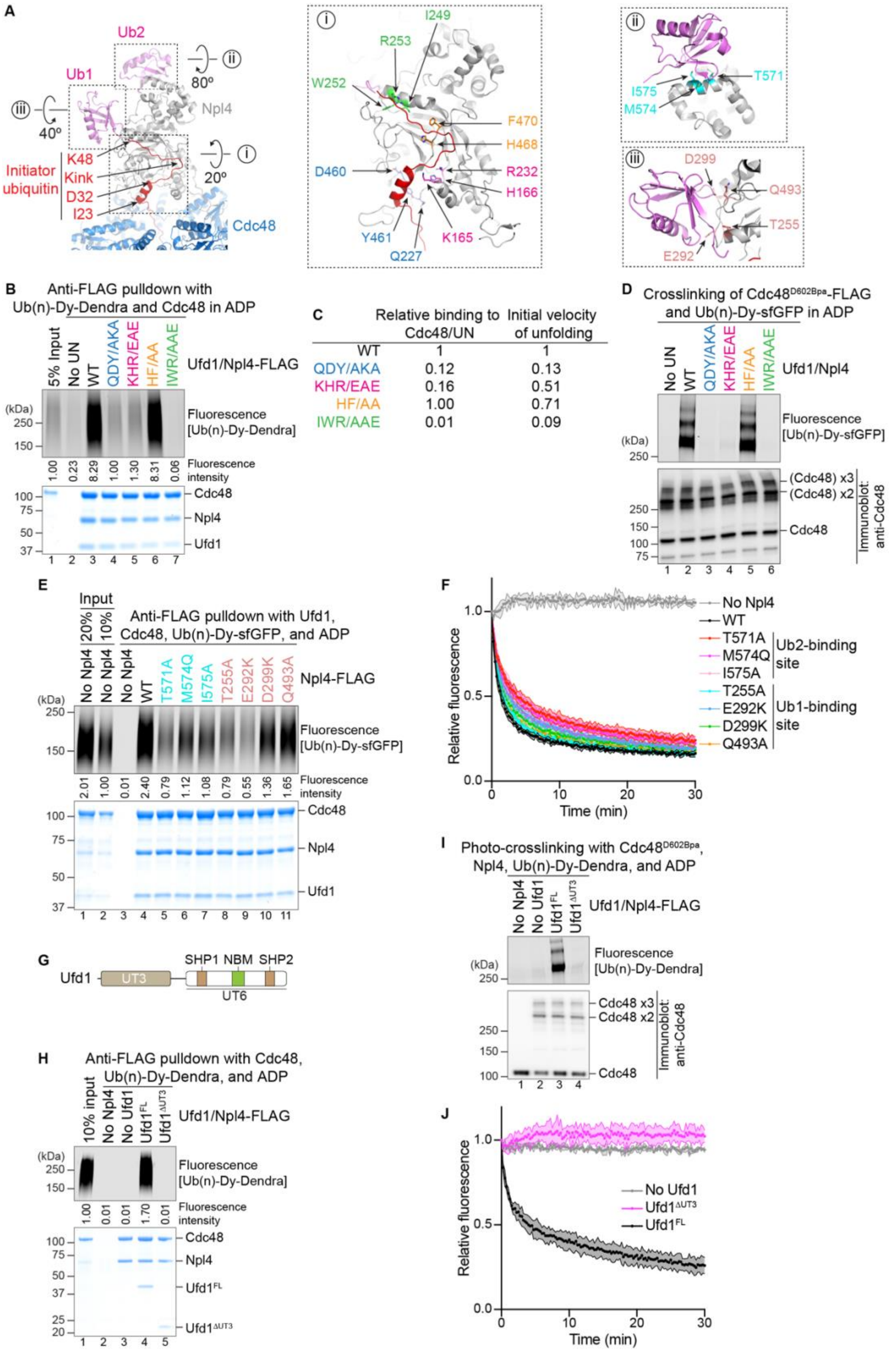


C

| | Relative binding to Cdc48/UN | Initial velocity of unfolding |
|---|---|---|
| WT | 1 | 1 |
| QDY/AKA | 0.12 | 0.13 |
| KHR/EAE | 0.16 | 0.51 |
| HF/AA | 1.00 | 0.71 |
| IWR/AAE | 0.01 | 0.09 |

**Figure S3. (related to Figure 3). Ubiquitin unfolding requires the Cdc48 cofactors Npl4 and Ufd1.**

**(A)** Model of Npl4 with bound ubiquitin molecules, based on a cryo-EM structure of a Cdc48-substrate complex (PDB, 6OA9). Npl4 is shown in grey, the unfolded initiator ubiquitin molecule in red, the two folded ubiquitin molecules Ub1 and Ub2 in pink, and Cdc48 in blue. The insets show magnified views with mutated residues highlighted in different colors. Residues shown in the same color in inset (i) were mutated simultaneously.

**(B)** Photoconverted dye-labeled Ub(n)-Dendra was incubated in the presence of ADP with Cdc48, Ufd1, and FLAG-tagged wild-type Npl4 (WT) or mutant Npl4 carrying the indicated mutations in the groove ((A); inset (i)). The Cdc48 complex was retrieved with FLAG antibody beads, and bound material analyzed by SDS-PAGE, followed by fluorescence scanning (upper panel) and Coomassie-blue staining (lower panel).

**(C)** Cdc48 complexes containing Npl4 mutants in the ubiquitin binding groove were tested for their binding of photoconverted dye-labeled Ub(n)-Dendra (in (B)) and their ability to unfold Dendra by measuring the initial rate of fluorescence loss. In both tests, wild-type (WT) Npl4 served as reference.

**(D)** Cdc48 complex containing FLAG-tagged Cdc48 with a photoreactive Bpa probe in the D2 domain at position D602 (D602Bpa) was incubated with dye-labeled Ub(n)-sfGFP, Ufd1, and the indicated Npl4 proteins. The samples were irradiated with UV light. Cdc48-containing complexes were retrieved with FLAG antibodies beads and analyzed by SDS-PAGE, followed by fluorescence scanning and immunoblotting with anti-Cdc48 antibodies. Note that the substrate not only crosslinks to monomeric Cdc48, but also to crosslinked dimers and trimers.

**(E)** As in (B), except that Npl4 residues interacting with Ub1 and Ub2 were mutated ((A); insets ii and iii).

**(F)** Cdc48 complexes containing wild-type (WT) Npl4 or Npl4 mutants at the binding interface to the folded ubiquitin molecules Ub1 and Ub2 were tested for unfolding of photoconverted Ub(n)-Dendra. The experiments were performed in triplicates. Shown are means and standard deviations for each data point.

**(G)** Domain structure of Ufd1. UT3 binds ubiquitin, SHP1 and SHP2 interact with Cdc48, and NBM binds to Npl4.

**(H)** As in (B), except that full-length Ufd1 ($Ufd1^{FL}$) and Ufd1 lacking the ubiquitin-binding UT3 domain ($Ufd1^{\Delta UT3}$) were compared in substrate binding experiments.

**(I)** As in (D), except that $Ufd1^{FL}$ and $Ufd1^{\Delta UT3}$ were compared in photocrosslinking experiments.

**(J)** As in (F), except that Dendra unfolding was tested with Cdc48 complex containing $Ufd1^{FL}$ or $Ufd1^{\Delta UT3}$.

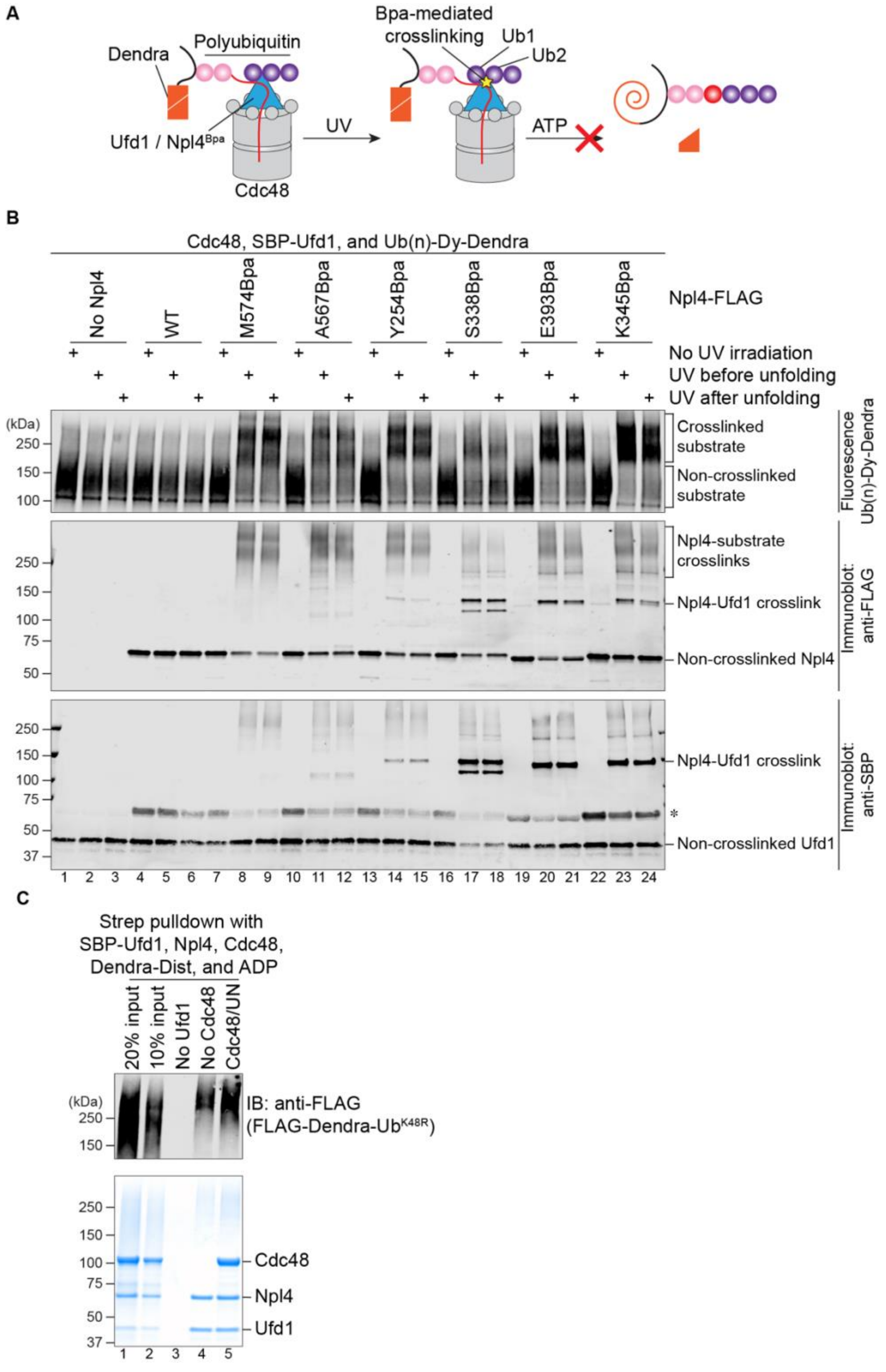


**Figure S4. (related to Figures 4 and 5). Site-specific photocrosslinking of Npl4 to bound ubiquitin molecules and binding of polyubiquitinated Dendra-Dist to Cdc48 complex.**

**(A)** Scheme for testing the translocation of the initiator ubiquitin. Photoreactive Bpa groups were incorporated into Npl4 at different positions, and crosslinks to either the unfolded initiator ubiquitin or the folded ubiquitin molecules Ub1 and Ub2 were induced by UV light. If Cdc48 begins substrate translocation by pulling on the initiator ubiquitin, crosslinking to Npl4 would prevent subsequent ATP-dependent Dendra unfolding.

**(B)** Photoconverted, dye-labeled Ub(n)-Dendra was incubated with Cdc48, SBP-Ufd1, and either wild-type (WT) Npl4 or Npl4 mutants carrying photoreactive Bpa probes at different positions (see **Figure 4A**). The samples were irradiated with UV light either before or after unfolding of Dendra in ATP. Where indicated, UV irradiation was omitted. All samples were analyzed by SDS-PAGE, followed by fluorescence scanning (upper panel) and immunoblotting with FLAG antibodies (detecting Npl4) and anti-SBP antibodies (detecting Ufd1) (lower two panels). Note that the anti-SBP antibodies cross-react with the Npl4 protein, as indicated by the star symbol.

**(C)** Dendra-Dist, generated by attaching FLAG-tagged Dendra-Ub$^{K48R}$ to the distal end of a K48-linked polyubiquitin chain (**Figure 5B**), was incubated with SBP-tagged Ufd1, Npl4, and Cdc48 in the presence of ADP. The Cdc48 complex was retrieved with streptavidin beads, and bound material analyzed by SDS-PAGE, followed by immunoblotting (IB) with FLAG antibodies (upper panel) and Coomassie-blue staining (lower panel).

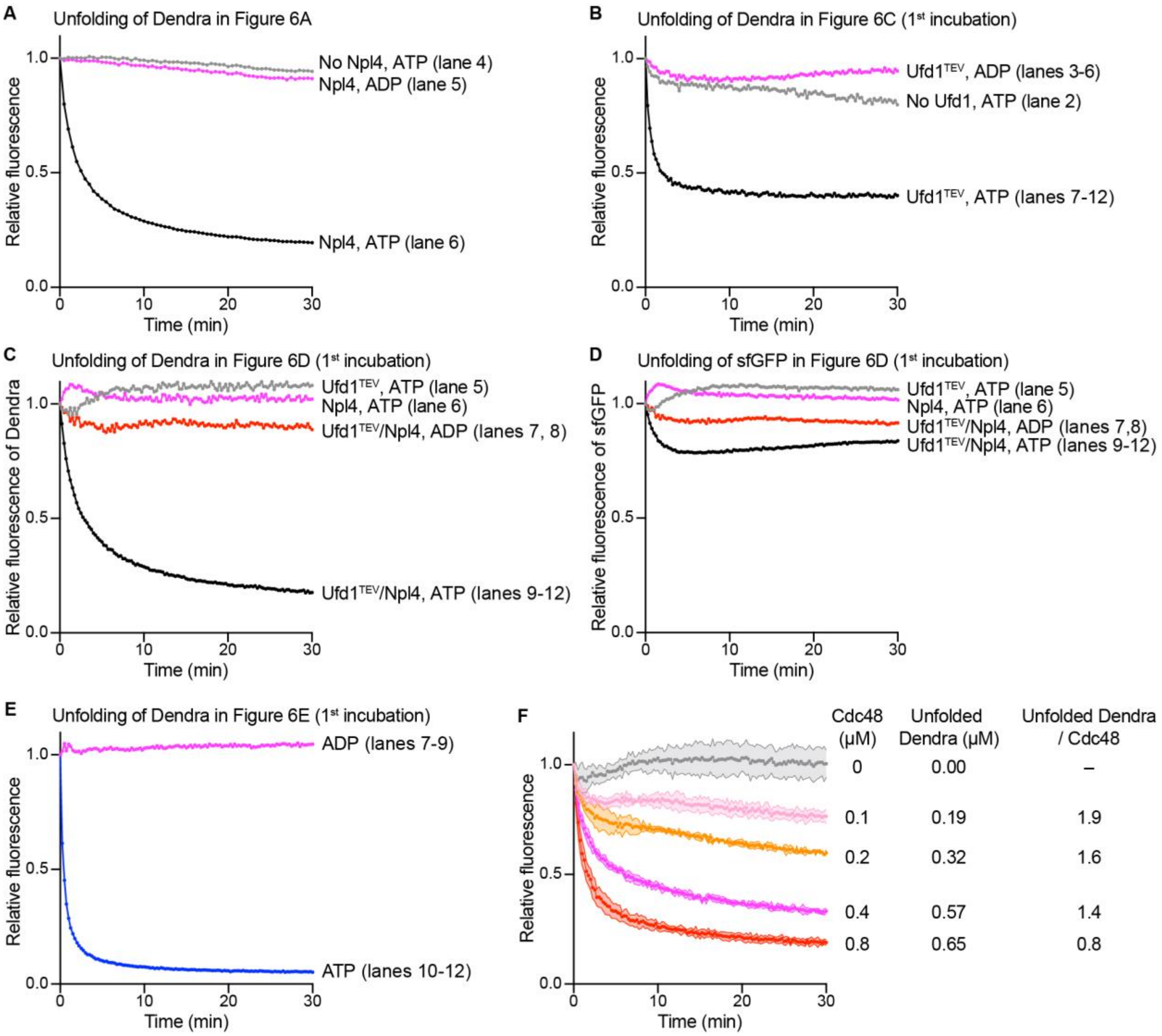


**Figure S5. (related to Figure 6). The Cdc48 complex releases unfolded, polyubiquitinated substrate and can translocate more than one substrate molecule.**

**(A)** The samples used in **Figure 6, lanes 4-6,** were analyzed for Dendra unfolding, as measured by the loss of fluorescence. Unfolding only occurs in the presence of Npl4 and ATP (lane 6).

**(B)** Samples in **Figure 6C** were analyzed during the 1$^{st}$ incubation in the absence of Ufd1 (lane 2) or in the presence of Ufd1$^{TEV}$ and either ADP (lanes 3-6) or ATP (lanes 7-12) for Dendra unfolding. Note that uncleaved Ufd1$^{TEV}$ supports Dendra unfolding.

**(C)** Samples in **Figure 6D** were analyzed during the 1$^{st}$ incubation in the absence of Npl4 (lane 5) or Ufd1 (lane 6) or in the presence of Ufd1$^{TEV}$ and Npl4 in either ADP (lanes 7,8) or ATP (lanes 9-

12) for Dendra unfolding. Dendra is irreversibly unfolded when all components and ATP are present.

**(D)** The samples in (C) were analyzed for sfGFP unfolding. Note the transient unfolding of sfGFP when all components and ATP are present (black curve).

**(E)** Samples in **Figure 6E** were analyzed during the 1$^{st}$ incubation in the presence of ADP (lanes 7-9) or ATP (lanes 10-12) for Dendra unfolding.

**(F)** Photoconverted Ub(n)-Dendra was incubated in ATP with UN and indicated amount of Cdc48. Unfolding of Dendra was followed (left panel) and the final level of unfolding after 30 min incubation was determined. These numbers were used to calculate how many substrate molecules were unfolded by a Cdc48 hexamer. The experiments were performed in triplicates. Shown are means and standard deviations for each data point.

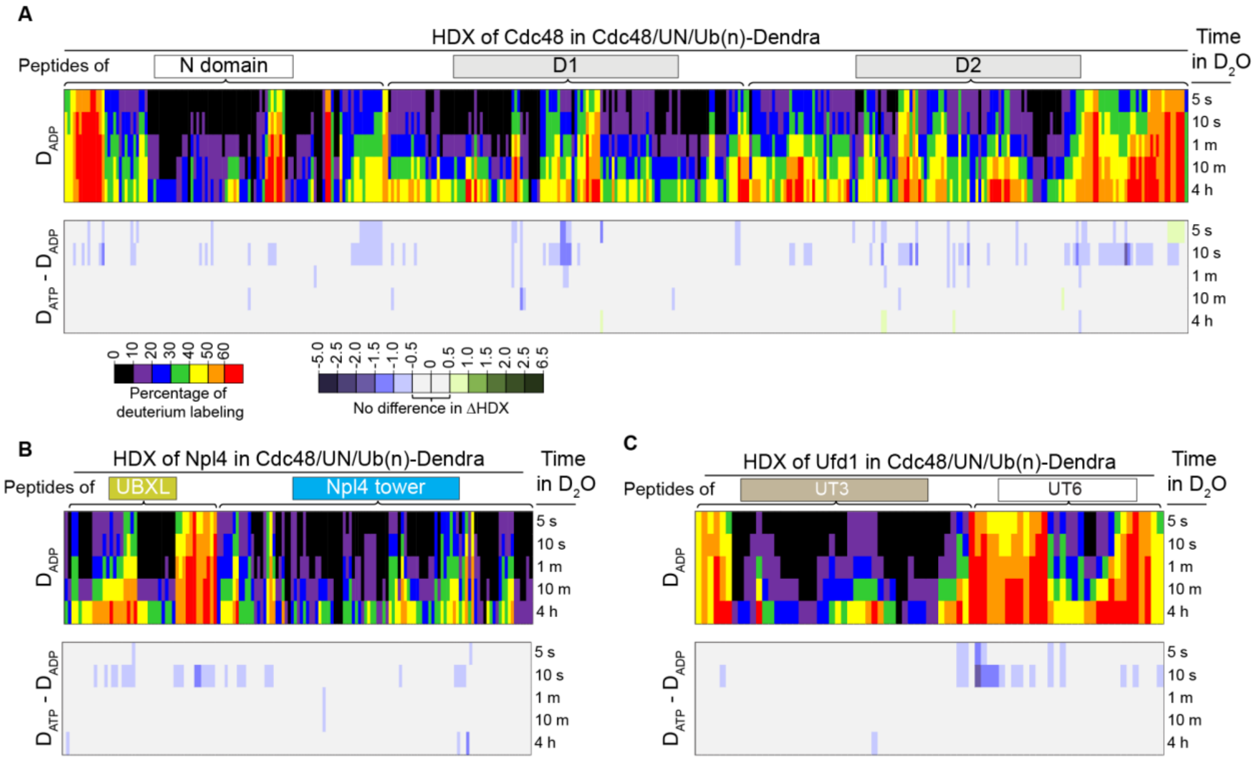


**Figure S6. (related to Figure 6). The Cdc48 complex is in a similar state before and after substrate unfolding.**

**(A)** Photoconverted Ub(n)-Dendra was incubated with Cdc48/UN complex in the presence of ADP or ATP. The samples were then subjected to HDX for different time periods as indicated, labeling was quenched, the proteins were subjected to proteolysis, and the deuteration of Cdc48 peptides was determined by MS. Peptides are arranged from N- to C-terminus left to right, and the position of the domains is indicated. The upper panel shows the relative percent deuteration of Cdc48 observed in ADP and the lower panel the difference in deuteration ($D_{ATP}$ – $D_{ADP}$) for Cdc48 in the presence of ATP versus ADP (scales given at the bottom). All peptide sequences and measured values used to create this figure are provided in the **Table S1**.

**(B)** As in (A), but for Npl4.

**(C)** As in (A), but for Ufd1.

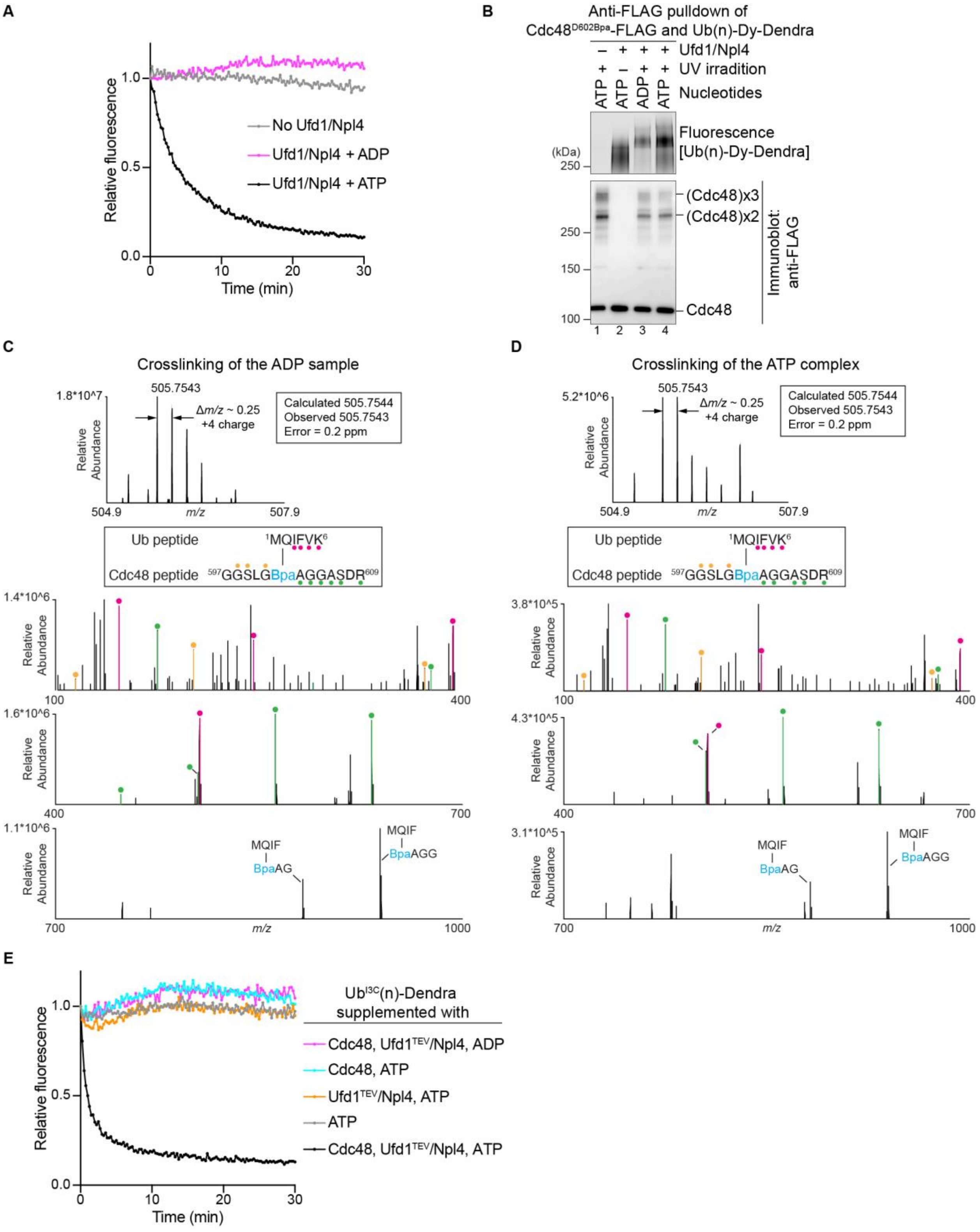


**Figure S7. (related to Figures 6 and 7). Polyubiquitinated substrate rebinds to the Cdc48 complex after unfolding.**

**(A)** Photoconverted dye-labeled Ub(n)-Dendra was incubated in the presence of ADP or ATP with FLAG-tagged Cdc48 complex containing a photoreactive Bpa probe in the D2 ring (position 602). The unfolding of Dendra was followed by the loss of fluorescence. After 30 min, the samples were used for crosslinking (panel B).
**(B)** The samples in (A) were irradiated with UV light, the Cdc48 complex was retrieved with FLAG antibody beads and analyzed by SDS-PAGE, followed by fluorescence scanning (upper panel) and immunoblotting with FLAG antibodies (lower panel).
**(C)** The ADP-treated sample shown in panel B (lane 3) was treated with trypsin and peptides analyzed by nano-liquid chromatography/MS. The upper panel shows the detection of a quadruply charged precursor ion at *m/z* 505.7543, consistent with the mass of the indicated crosslinked Cdc48 and ubiquitin peptides. The lower panel shows the MS/MS spectrum of this crosslinked peptide precursor. Cdc48 derived ions of type b and y are marked with yellow and green glyphs, respectively, while y-type ions derived from ubiquitin are marked with magenta glyphs. Ions containing the crosslinked peptides were also detected in the MS/MS spectrum.
**(D)** As in (C), but for the ATP-treated sample shown in panel B (lane 4).
**(E)** Photoconverted $Ub^{I3C}$(n)-Dendra was incubated in the presence of ADP or ATP with different combinations of Cdc48, Npl4, and Ufd1 containing a TEV protease cleavage site following the UT3 domain ($Ufd1^{TEV}$). The unfolding of Dendra was followed by the loss of fluorescence.
